# Ubiquitin-Proteasome System Dysregulation in Alzheimer’s Disease Impacts Protein Abundance

**DOI:** 10.1101/2025.05.29.656728

**Authors:** Mahlon Collins, Corinna Friedrich, Megan Elcheikhali, Peyton Stewart, Jason Derks, Theresa Connors-Stewart, Kirsten Altig, Alexandra Melloni, Aleksandra Petelski, Derek Oakley, Bradley Hyman, Nikolai Slavov

**Affiliations:** Parallel Squared Technology Institute, Watertown, MA, USA; Department of Neurology, Massachusetts General Hospital, Charlestown, MA, USA; Department of Pathology, Massachusetts General Hospital, Charlestown, MA, USA; Department of Bioengineering, Northeastern University, Boston, MA, USA

## Abstract

Alzheimer’s disease (AD) is a relentlessly progressive, fatal neurodegenerative disorder that results in widespread protein dysfunctions. However, the full extent of aberrant proteomic changes in AD and their impact remains unknown, in part, because of the challenges of comprehensively measuring the proteome. Here, we used plexDIA, an approach that provides deep proteomic coverage and high throughput by parallelizing the acquisition of peptides and samples, to characterize proteomic changes in AD. Using human dorsolateral prefrontal cortex tissue, we identified 281 differentially abundant proteins in AD. By systematically analyzing compartment and protein complex-specific shifts in protein abundance, we identified an AD-specific decrease in levels of the 20S proteasome, the catalytic core of the cell’s primary protein degradation pathway. This alteration was accompanied by widespread decreases in proteasome subunit stoichiometries. Many proteasome substrate proteins were negatively correlated with 20S levels and increased in AD, suggesting that reduced 20S levels leads to abnormal protein accumulation. By analyzing proteins increased in AD, we identify key properties of such proteins: They have fast degradation rates, they contain signal sequences that allow them to be targeted for proteasomal degradation, and they are targeted by quality control pathways that recognize mislocalized proteins. Changes in these gene products at the protein and mRNA levels were highly discordant, providing additional evidence for increased protein abundance driven by impaired clearance. We also identified coherent sets of ubiquitin system enzymes, proteins that target substrates for proteasomal degradation, whose levels robustly discriminate AD from non-AD samples. One subset exhibited consistent increases in AD, while another, which contained the tau E3 ligase Cul5, exhibited consistent decreases, revealing complex changes in the ubiquitin system in AD. Taken together, our results suggest that decreased ubiquitin-proteasome system capacity and impaired clearance of short-lived and mislocalized proteins contribute substantially to proteopathic burden in AD.

## Introduction

Alzheimer’s disease (AD), the most common form of dementia, results in progressive memory loss, emotional disturbances, and cognitive dysfunction^1,2^. The disease’s defining pathological features are filamentous intracellular inclusions containing hyperphosphorylated tau and amorphous extracellular aggregates of amyloid beta peptide^3–5^. However, AD is increasingly recognized as a multi-proteopathy with protein dysfunctions affecting a multitude of proteins with distinct sequence compositions, structures, functions, and subcellular localizations. For example, neuronal inclusions containing the synaptic protein alpha-synuclein^6,7^, the nuclear RNA-binding protein TDP-43^6,8^, and components of the U1 small nuclear ribonucleoprotein spliceosome^9,10^ often occur in AD-afflicted neurons. These protein lesions exert numerous downstream effects on neuronal physiology and protein homeostasis^11–13^ which may further exacerbate the development and spread of AD protein pathologies and neuron loss^14^. The full extent of protein dysfunctions in AD and the mechanisms that give rise to them remain incompletely characterized, limiting our understanding of the disease’s causes and hindering efforts to develop disease-modifying therapies^15^.

Mass spectrometry (MS) provides a comprehensive approach for characterizing AD-associated protein dysfunctions. MS instruments can directly measure a sample’s complement of proteins with high sensitivity and quantitative accuracy^16,17^. Using such data to identify individual proteins with large fold changes between AD and non-AD samples can reveal novel AD protein dysfunctions and pathologies^9,10^ in addition to robustly detecting increases in proteins known to form pathological aggregates in AD^9,10,18,19^. AD protein pathologies reflect a failure of quality control mechanisms to clear aberrantly misfolded, damaged, and aggregated proteins from cells^12,13^. As a post-mitotic cell population, neurons are acutely vulnerable to these challenges, as they cannot reduce proteopathic burden by dilution through cell division^13,20^. However, because of the large changes in protein abundance that result from AD protein pathologies, these changes can be readily detected in bulk brain tissue samples containing multiple cell types with varying degrees of AD pathology^9,10,18,19,21,22^. Proteomic analysis of bulk samples thus holds considerable promise for identifying novel AD protein dysfunctions and pathologies.

Proteomic data has also proven valuable for providing systems-level insights into the molecular mechanisms of AD and AD-associated protein pathologies. Pairing large-scale proteomic data with systems-level analytical approaches can detect alterations to biological processes and cellular compartments in AD^23–25^. For example, proteomic profiling of AD samples detects the expected strong and significant downward shift in cytoskeletal and synaptic proteins, reflecting the loss of cytoskeletal integrity following tau dissociation from microtubules and synaptic dysfunction, respectively^23–25^. Upward shifts in inflammation-related proteins similarly reflect glial activation in AD^26^. Network analysis is a related approach that identifies disease modules, clusters of highly correlated proteins altered in disease^10,19,27^. Proteomic data has been used to identify dozens of AD modules, highlighting altered signaling networks, metabolic pathways, and cell states in AD^23–25, 28^.

Although MS proteomics has been productively used to identify novel individual AD protein pathologies and network alterations, the approach’s full potential has yet to be realized in the context of AD. In particular, sample throughput, defined as the number of parallel samples and proteins that can be analyzed, remains limiting. Approaches that analyze a single sample per MS run (“label-free”) provide excellent quantitative accuracy and identify many proteins^16,29^. However, AD is a highly heterogeneous disease influenced by complex genetic and lifestyle factors^15,30–32^. Understanding the disease’s causal mechanism thus requires profiling large cohorts, which is impractical with label-free approaches. Molecular barcodes (“mass tags”) allow multiple samples to be pooled and run simultaneously^33^. However, multiplexing samples often results in fewer proteins identified per sample and quantification relying on isobaric mass tags is often adversely affected by co-isolation interfer-ence^34,35^. Further, the vast majority of prior AD MS proteomic profiling efforts have used data-dependent acquisition, an approach that isolates and fragments one peptide precursor at a time. In an alternative framework, data-independent acquisition (DIA), all precursors within a specified window are analyzed, increasing throughput and data completeness by parallelizing the analysis of peptides^36,37^. Recent advances have combined non-isobaric (differing mass) mass tags with DIA to increase throughput in MS proteomics^38,39^. The resulting experimental and computational framework, plexDIA, provides multiplicative gains in throughput by simultaneously multiplexing samples and peptides^38,39^. Quantification with plex-DIA is based on peptide-specific ions^38,40^, so it is not affected by the co-isolation interference that undermines the accuracy of TMT-based multiplexing approaches.

Here, we leveraged plexDIA to quantify proteins in human AD and non-AD brain tissue samples. We used the resulting datasets to identify individual protein alterations and systematic changes in proteins mapped to well-annotated biological processes, cellular compartments, and protein complexes. The largest systemic shift in our data was an AD-specific reduction in the 20S proteasome, the catalytic core of the primary protein degradation pathway in eukaryotic cells. This reduction coincided with reduced subunit stoichiometry both within and between the proteasome’s two functional modules: the 20S core particle and 19S regulatory particle. Analyzing the relationship between 20S levels and protein abundance revealed unique properties of proteins that increase AD. First, they tend to have faster degradation rates than those decreased in AD. Second, proteins increased in AD harbor more signals that enable them to be targeted for degradation by the ubiquitin-proteasome system (UPS). Third, a number of proteins with large increases in AD have compartment-specific subcellular localizations and are targeted for UPS degradation when mislocalized. Taken together, our results are consistent with a model in which both decreased ubiquitin-proteasome system capacity and aberrant protein accumulation adversely impact protein homeostasis in AD.

## Results

### Widespread Proteomic Changes in AD Identified using plexDIA

We sought to leverage plexDIA’s high data completeness and throughput^38,41^ to characterize AD proteomic changes. To do so, we acquired brain tissue samples from a cohort of 24 individuals from the Massachusetts Alzheimer’s Disease Research Center (ADRC). Tissue from Brodmann area 9 of the dorsolateral prefrontal cortex was used for all analyses. Hallmark AD tau pathology occurs primarily in non-cortical structures in Braak stage III or lower cases^5,42,43^. To increase our power to see AD-related proteomic changes, we therefore categorized subjects at Braak stages 0-III “non-AD” and V and VI as “AD” (Figure 1A). We labeled individual samples with the non-isobaric (differing mass) mTRAQ labeling reagents, which enables 3-plex multiplexing. This resulted in 8 sample batches each with similar distributions of age, sex, and disease stage (Figure 1A; Methods). To increase the specificity of sequence identification, we created a sample-specific spectral library using narrow isolation windows that allowed for high specificity mapping between precursors and their corresponding fragment ions^44^. This allowed for confident identification of peptides and their modification and the creation of specific spectral libraries for searching all spectra. Using a Thermo Exploris 480 instrument and DIA-NN^41^, we confidently identified and quantified 6,436 proteins across the set of 24 samples. To understand to what extent proteomic changes were reflected in the transcriptome, we also performed RNA-seq on the same set of samples.

**Figure 1:**
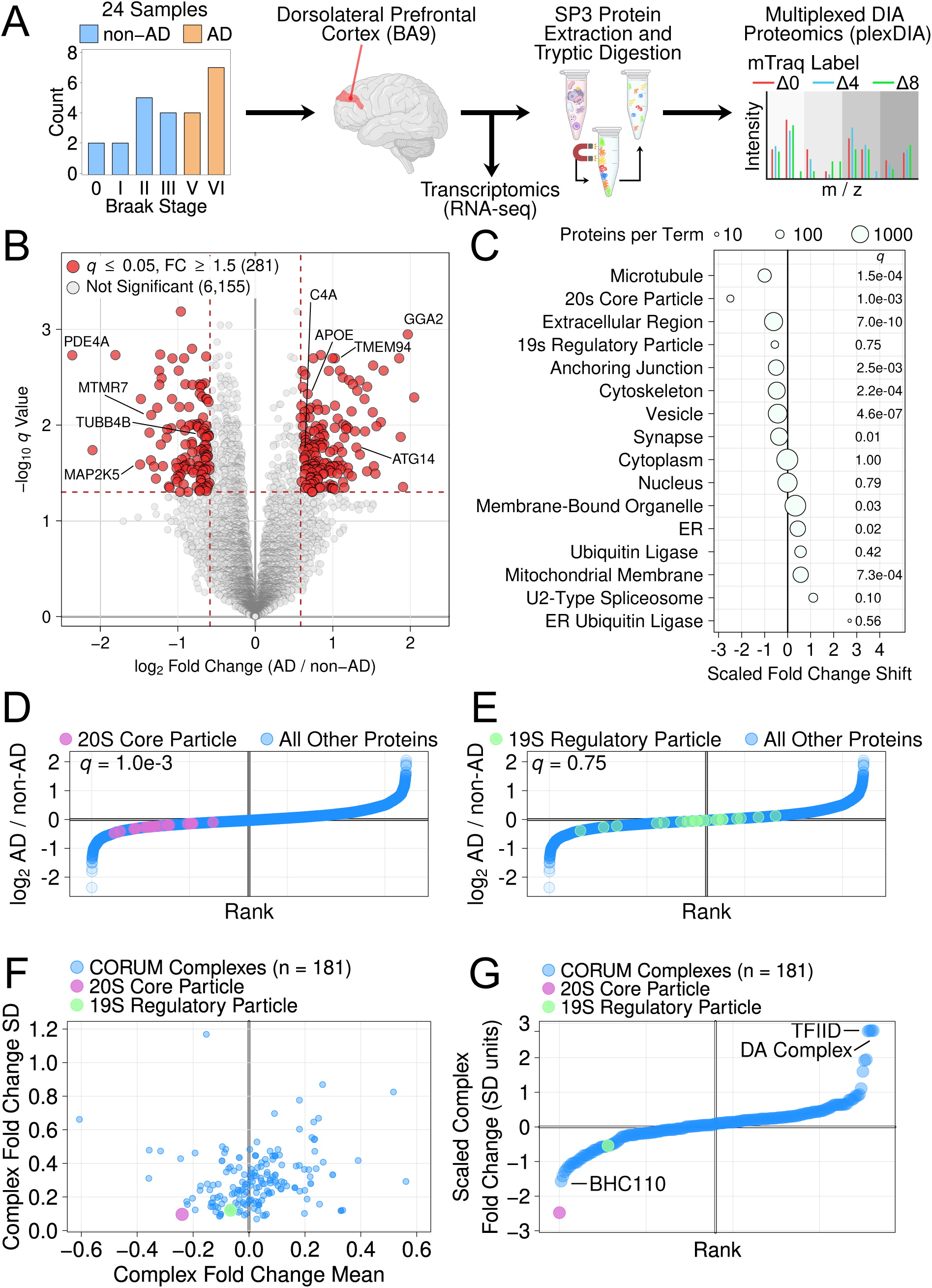
Study overview and AD proteomic changes. **A**. Schematic of the cohort and experimental approaches. **B**. Volcano plot of proteins exhibiting differential abundance (q value less than 0.05 and an absolute fold change greater than 1.5) between AD and non-AD samples. **C**. Compartment-specific shifts were scaled and plotted to visualize the largest shifts between AD and non-AD, revealing that the shift in proteasome 20S subunit proteins was the largest negative shift between AD and non-AD samples. **D.** / **E.** Rank order plots showing relative fold change ranks of 20S (D.) and 19S (E.) proteasome subunits between AD and non-AD samples. **F.** The mean shift of 183 protein complexes with at least 5 subunits detected in our data in the CORUM database is plotted against the standard deviation. **G.** Scaled protein complex shifts were computed by dividing a complex’s mean shift by its standard deviation and visualized as a rank-order plot, revealing that the largest decrease in AD was that of 20S core particle levels.

Exploratory analyses revealed that our data accurately captured known AD biology. We first visualized samples using principal component analysis (PCA), which revealed that samples separated clearly on the basis of disease status (“AD” or “non-AD”) along the first principal component (Supplementary Figure 1). Samples did not obviously separate on the basis of other demographic or technical factors unrelated to disease status (Supplementary Figure 1). We then determined the set of differentially abundant proteins between AD and non-AD samples. To identify robust, consistent signals in our data, we used a significance criteria of a *q* value cutoff of 0.05 and an absolute AD / non-AD fold change greater than 1.5. In total, 281 proteins met these criteria. (Figure 1B). The set of differentially abundant proteins included many proteins with known roles in AD, such as increased APOE^18,21,45^ and the inflammation-related complement C4A^18,21,46^, as well as decreased levels of cytoskeletal proteins^18,21,47^ (MAP4 and tubulin beta-4B chain) and mitogen-activated protein kinase kinase 5 (MAP2K5) (Figure 1B). The largest absolute fold change was the significant decrease in the cAMP phosphodiesterase PDE4A, a change previously captured in large-scale studies of AD-associated proteomic changes^18,21^ and a potential AD therapeutic target^48^.

AD is associated with widespread alterations in multiple cellular organelles, compartments, and complexes, including the neuronal cytoskeleton^47,49^, the endolysosomal system^11,50^, and the endoplasmic reticulum (ER)^51,52^. Although changes in selected individual proteins have been well-characterized^53–55^, a comprehensive compartment-specific census of AD-linked proteomic changes has not been achieved.

To this end, we applied a recently-described approach for identifying shifts in specific cellular organelles, compartments, and complexes from bulk proteomics data^56^. The approach searches for systematic shifts in fold changes of proteins annotated to specific cellular compartments. To assess the approach in the context of AD, we first tested whether proteins annotated to the cytoskeleton exhibited consistent shifts between AD and non-AD samples. As expected, we observed a large, statistically significant (Wilcoxon *q* = 2.2e-4) shift, such that levels of cytoskeletal proteins were consistently lower in AD (Figure 1C). We obtained similar results for microtubule proteins (Wilcoxon *q* = 1.5e-4).

Having established the approach’s ability to capture known AD compartment-specific proteomic changes, we next applied it across the set of Gene Ontology^57,58^ Cellular Component terms. The largest AD-associated shift was the decrease in proteasome 20S core particle subunits (Wilcoxon *q* = 1e-3; Figure 1C / D), consistent with the reduced proteasomal capacity recently observed in other human AD samples^59–61^. The 20S is the degradative core of the 26S proteasome, the primary protein degradation pathway in eukaryotic organisms^62–64^. The canonical configuration of the proteasome, the 26S form, consists of a single 20S core particle doubly capped with 19S regulatory particles. In this configuration, the 19S regulatory particle binds ubiquitinated substrates and processively unfolds and translocates them to the 20S core, where substrates are degraded to short peptides^63–66^. Emerging evidence also suggests that free 20S core particles are abundant within cells and exhibit distinct substrate preferences as compared to 26S proteasomes^67–69^. Thus, the shift in 20S core particles in AD may reflect decreased degradative capacity in AD-afflicted cells, further compounding proteostatic challenges induced by AD protein pathology^68,70,71^. We applied the same analysis to the 19S regulatory particle. Although selected subunits exhibited a leftward shift, the overall change was non-significant (Wilcoxon *q* = 0.75; Figure 1C / E).

To put these results in context, we scaled all compartment-specific fold change shifts relative to that observed for microtubules, a large, significant change that reflects known AD biology. Doing so revealed multiple compartment-specific shifts. Some shifts likely reflect neuronal loss in AD, such as decreases in synaptic (*q* = 0.01) and vesicle-associated proteins (*q* = 4.6e-7). Others likely reflect impaired proteostasis within AD-afflicted cells, such as increased levels of ER proteins (*q* = 0.02; Figure 2C). The largest increase for AD was observed for ER ubiquitin ligases (Figure 2C), however this term did not reach statistical significance, likely owing to the small number of associated proteins, six, detected in our study. Taken together, our results reveal proteomic changes across a diverse range of organelles, compartments, and complexes in AD.

**Figure 2:**
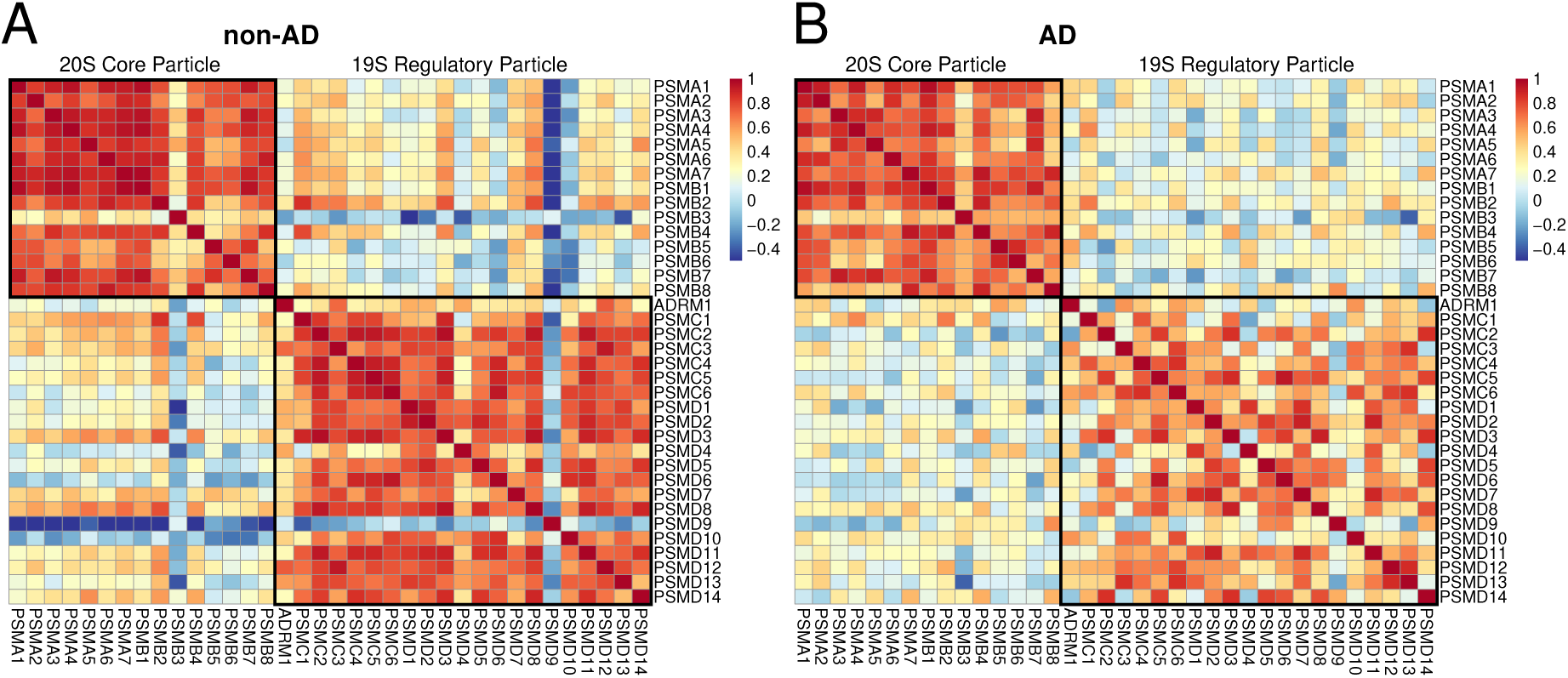
Proteasome subunit stoichiometries in non-AD and AD samples. Correlations between the abundance of individual proteasome subunits were visualized as a heatmap for non-AD and AD samples. **A. / B.** non-AD samples (A.) showed significantly higher correlations among components of the 19S regulatory particle and 20S core particle as compared to AD samples (B.).

To provide additional context to our results, we assessed how the change in 20S proteasome levels in AD compared to other multimeric protein complexes. For this, we used the set of complexes in the CORUM database^72,73^. We restricted our analysis to the 183 complexes with at least 5 subunits that were detected in our data. Visualizing results for these complexes revealed that the decrease in 20S levels was among the largest and most consistent decreases among protein complex (Figure 2F). To provide a scaled measure of the shift in complex abundance, we divided the mean fold change shift of all complex subunits by their standard deviation. By this scaled metric, the shift in 20S levels was the largest decrease among protein complexes in AD (Figure 2F). The next most decreased complex was BHC110, a corepressor complex involved in transcriptional silencing. Several complexes also showed large increases in AD. Among these, the largest shifts were observed for the DA and TFIID complexes, complexes involved in RNA polymerase II (Pol II)-dependent transcription. These complexes contain highly overlapping sets of gene products, in particular, the RNA polymerase II, TATA box binding protein-associated factors (TAFs). The increased levels of various TAF proteins in AD observed here are consistent with prior results^74,75^. Taken together, our results reveal that the decreased 20S levels are the largest change among both annotated subcellular compartments and protein complexes in our cohort.

### Reduced Proteasome Subunit Stoichiometry in AD

The 20S proteasome consists of a barrel-shaped arrangement of stacked *α* and *β* subunits arranged as heteroheptameric rings^76–78^. Consistent with these structural requirements, the production rate of 20S subunits is generally highly similar between subunits^79^ and the abundance of individual subunits is highly correlated^80,81^. In contrast, the 19S regulatory particle can exist in multiple configurations and multiple subunits are often produced in stoichiometric excess^82–85^. To understand if the shift in 20S core particle abundance reflects a reduced stoichiometry among 19S and 20S complexes, we computed correlations within and between individual proteasome subunits of the 19S regulatory particle and 20S core particle. As expected, the correlation among 20S core particles subunits was large and positive. However, AD samples showed significantly reduced correlations among all 20S subunits (Wilcoxon *p* = 0.021; Figure 2).

An even more marked reduction in subunit correlations was observed for the 19S regulatory particle in AD cases (Wilcoxon *p* = 3e-17; Figure 2). Together, these results reveal a widespread decoupling of the abundances of the subunits of the proteasome that may result from or further exacerbate defects in protein quality control pathways in AD^11,59–61,71^.

### Multiple AD-relevant Protein Sets are Correlated with 20S Levels

To better understand the relationship between proteasome levels and protein abundance, we correlated levels of individual proteasome subunits with all other proteins across our 24 samples. Owing to their highly similar abundances, protein correlations were consistent among 20S subunits (Figure 3A). We observed three distinct bands, one with mostly positive, one with mostly negative, and one with low overall correlation to 20S levels (Figure 3A, left to right). In contrast, proteome-wide correlations to 19S regulatory particle levels displayed two distinct clusters and much lower overall consistency among subunits (Figure 3B). We reasoned that we could use the median of 20S correlations to individual proteins to identify categories of proteins well-correlated to 20S levels. Multiple proteins with known roles in AD positively correlated to 20S levels. HSPE1 is a chaperone protein involved in the mitochondrial unfolded protein response^86^ and also implicated in AD^87^. Consistent with their roles in protein quality control, HSPE1 and 20S levels were significantly positively correlated (*r* = 0.67, *q* = 0.03). Variation in the *NTM* gene encoding neurotrimin, a neural cell adhesion molecule, modulates tau pathology levels in AD^88^. Neurotrimin levels were positively correlated with 20S levels (*r* = 0.74, *q* = 0.014). Calsystenin-1 (CLSTN1) is involved in axonal transport of amyloid beta and the protein is reduced in AD^89^. Calsystenin-1 levels were positively correlated with 20S levels (*r* = 0.64, *q* = 0.046). Reduced calsystenin-1 is known to trigger amyloid beta accumulation^89^, which likely exacerbates proteotoxic stress resulting from decreased proteasome capacity in AD.

**Figure 3:**
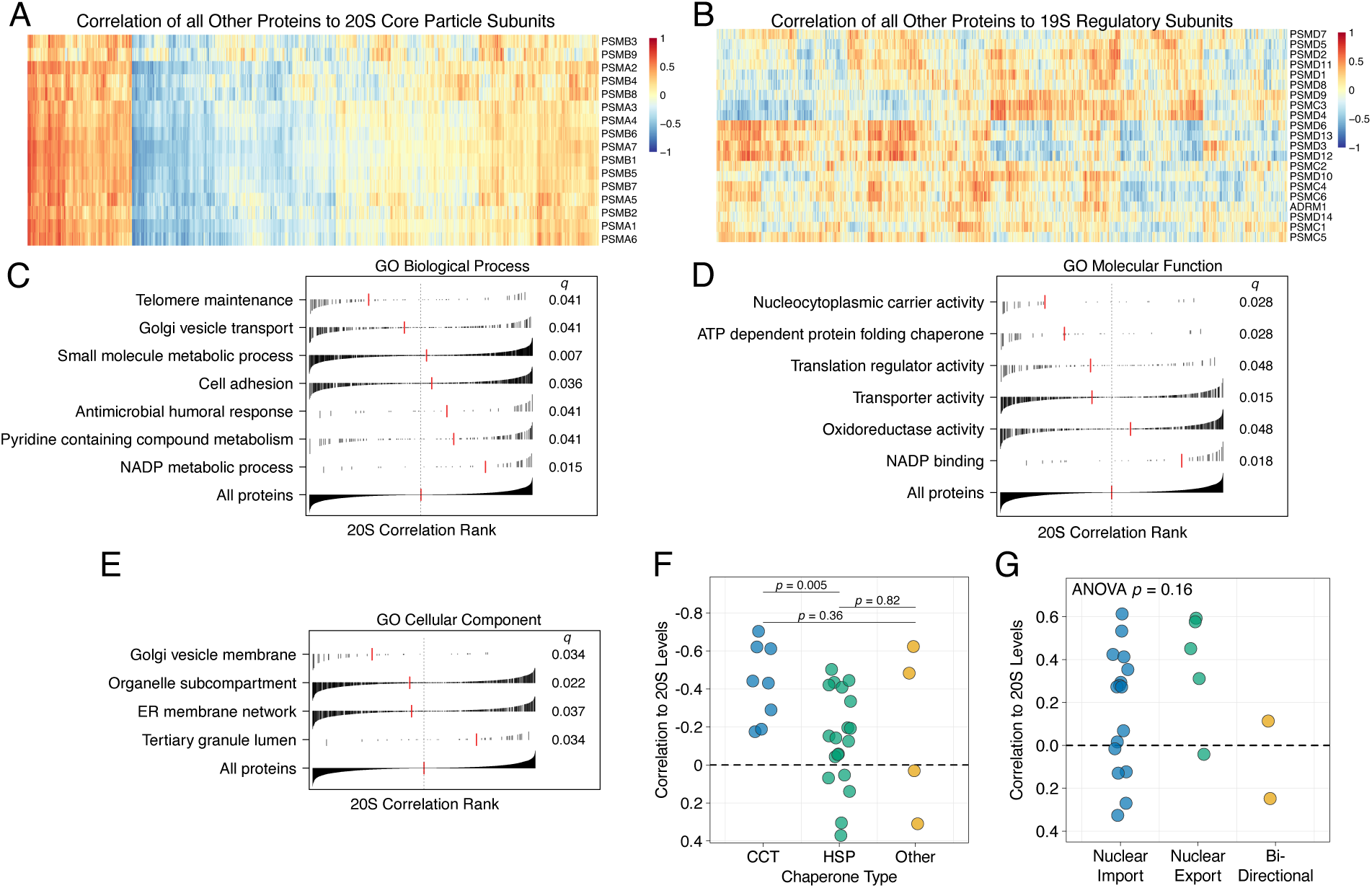
Proteome-wide correlations to 20S levels. **A.** Proteome-wide correlations to 20S proteasome core particle subunit levels. **B.** Proteome-wide correlations to 19S proteasome regulatory particle levels. **C. – E.** Levels of all proteins were correlated to 20S subunit median abundance and protein set enrichment analysis was run on the resulting set of correlations. Plots display significantly enriched (q ≤ 0.05) Gene Ontology Biological Process (C.), Molecular Function (D.), and Cellular Compartment (E.) terms. Black lines show individual proteins with the correlation magnitude displayed on the y axis. Red lines show the median for all proteins mapping to the indicated term. The full set of 20S correlations is shown for reference at the bottom of each plot in C - E. **F.** Proteins from the “ATP dependent protein folding chaperone” GO term were stratified by chaperone type and each protein’s correlation to 20S levels was visualized. As shown, proteins of the chaperonin containing TCP-1 (CCT) complex were significantly more negatively correlated to 20S levels than HSP chaperones. **G.** Proteins from the GO “Nucleocytoplasmic carrier activity” term were stratified by their direction of transport. There were no significant differences in 20S correlations between transporter types.

Proteins negatively correlated to 20S levels, which increase with decreasing 20S abundance, are of particular interest, since they likely include proteasome substrates. Correlation analysis alone cannot determine whether a protein rises to increased abundance due to reduced proteasome levels or activity. However, analyzing correlations and additional properties, such as degradation rate, number of degradation-targeting signals (“degrons”^90^), and subcellular localization can be used to generate testable hypotheses about protein homeostasis in AD. Ribosomal proteins accumulate in protein aggregates in the aging brain due to impaired clearance mechanisms, including reduced proteasome activity^91^. Consistent with this observation, we observed significant negative correlations between 20S levels and the ribosomal subunits RPS2 and RPL11 (*r* = −0.71, −0.66, *q* = 0.016 and 0.041, respectively). Excess synaptojanin 1 (a phosphoinositide phosphatase) contributes to synaptic defects in AD^92^. We observed a significant negative correlation of the protein to 20S levels (*r* = −0.8, *q* = 0.01). Aberrant increases in dynamin 1-like protein lead to mitochondrial defects and the protein has previously been implicated in AD^93^. We observed a significant negative correlation to 20S levels (*r* = −0.69, *q* = 0.02). Together, these analyses reveal multiple proteins with known connections to AD and strong correlations to 20S levels.

To provide a compartment- and function-specific view of proteomic correlations to 20S levels, we performed protein set enrichment analysis using the set of Gene Ontology (GO)^57,58^ Cellular Component, Biological Process, and Molecular Function terms. We identified 17 terms across the three domains at a 5% false discovery rate (FDR; Figure 3C-E). Overall, 9 of the 17 significant terms were driven by negative protein correlations to 20S levels. Protein localization to subcellular compartments or multi-protein complexes was a theme across the set of proteins negatively correlated to 20S levels, reflected at a high level in the significant enrichment for the GO Cellular Compartment term “Organelle subcompartment” (*q* = 0.022; Figure 3B). Specific compartments reflecting this enrichment included “Golgi vesicle membrane” (*q* = 0.034), “ER membrane network” (*q* = 0.037), “Nucleocytoplasmic carrier activity” (*q* = 0.028), and “translation regulator activity” (*q* = 0.048).

The main theme amongst enriched terms resulting from positive correlations to 20S levels was NADP metabolism. This was reflected in significant enrichments in the GO Biological Process terms “Pyridine containing compound metabolism” (*q* = 0.041) and “NADP metabolic process” (*q* = 0.015), as well as the molecular function terms “Oxidoreductase activity” (*q* = 0.048) and “NADP binding” (*q* = 0.018). There are no obvious functional or structural links between proteins from these terms and the proteasome. However, the significant positive correlation among these proteins may reflect similar energetic demands of ATP-dependent proteasomal protein degradation and NADP-dependent biosynthetic reactions. Thus, the observed positive correlations may reflect energetic imbalances, a well-characterized hallmark of AD^94–97^.

We next examined the set of proteins contained within selected significantly enriched terms. ATP-dependent folding chaperone proteins were negatively correlated with 20S levels and are elevated in AD, where they may mitigate proteotoxic stress and protein aggregation^98,99^. The chaperones contained in the “ATP-dependent folding chaperone” term comprise three main categories: those of the Chaperonin Containing TCP-1 (CCT) complex, heat shock proteins (HSPs), and other non-CCT / non-HSP chaperones. We stratified the set of proteins in our data based on these categories to determine whether a specific class was more strongly associated with 20S levels. CCT proteins were significantly more negatively correlated to 20S levels than HSP proteins (t-test *p* = 5e-3; Figure 3F). No other differences were observed among chaperone categories (t-test *p >* 0.05; Figure 3F). CCT members bind and potently inhibit tau aggregation^100^, providing a potential explanation for their robust induction in response to decreased 20S levels.

Nucleocytoplasmic transport defects have previously been described in AD and individual nuclear pore complex subunits accumulate in AD-afflicted neurons^101^. Similarly, we observed that nucleocytoplasmic transport proteins were significantly negatively correlated to 20S levels. The set of proteins contained in the GO “Nucleocytoplasmic carrier activity” term comprise proteins that facilitate nuclear import, nuclear export, or bi-directional transport of proteins between the nucleus and cytoplasm. We stratified proteins based on these categories to understand if a particular transport direction was more strongly associated with 20S levels. None of categories differed significantly in their correlation to 20S levels (ANOVA *p* = 0.16; Figure 3G).

### Increased Mitochondrial Ribosome Subunits Suggest Quality Control Defects in AD

To better understand our observation that many proteins targeted to specific organelles, subcellular compartments, or multi-protein complexes were inversely correlated with 20S levels, we examined proteins in our dataset that are known 20S substrates. 20S proteasomes are known to be highly abundant within cells and capable of directly degrading substrates without ubiquitination^62,67,68,102^. 20S proteasomes target distinct sets of substrates from 26S proteasomes. In particular, 20S proteasomes exhibit a high affinity for proteins with intrinsically disordered regions, presumably because 20S proteasomes lack the 19S regulatory particle’s unfolding capabilities^67,69,103^.

We identified 20S substrates in our data using previously-published datasets^69,104^. Examining the log_2_ fold change (AD / non-AD) of 20S substrates revealed that 20S substrates were twice as likely to be increased versus decreased in AD (Figure 4A). Among the set of significantly increased 20S substrates, MRPL54, a subunit of the mitochondrial ribosome large subunit, was a clear outlier, both in terms of absolute fold-change magnitude and *q* value (Figure 4A). Another mitochondrial 20S substrate, frataxin, was also significantly increased in AD (Figure 4A). PDCD4, a translational inhibitor^105^, was also significantly increased in AD. PDCD4 has long been known to be upregulated in AD^106^, but the mechanism by which this occurs is unknown. Our results raise the possibility that PDCD4 levels are increased, in part, as a result of decreased 20S proteasome levels. NUFIP2 is an intrinsically disordered RNA-binding protein that associates with cytoplasmic stress granules and shuttles between the nucleus and cytoplasm^107^. NUFIP2’s interaction with TDP-43 drives TDP-43 mis-localization and aggregation in a frontotemporal lobar degeneration (FTLD) model system^108^. The significantly increased levels of NU-FIP2 we observe may similarly contribute to TDP-43 dysfunction in AD through a failure of 20S proteasomes to degrade NUFIP2.

**Figure 4:**
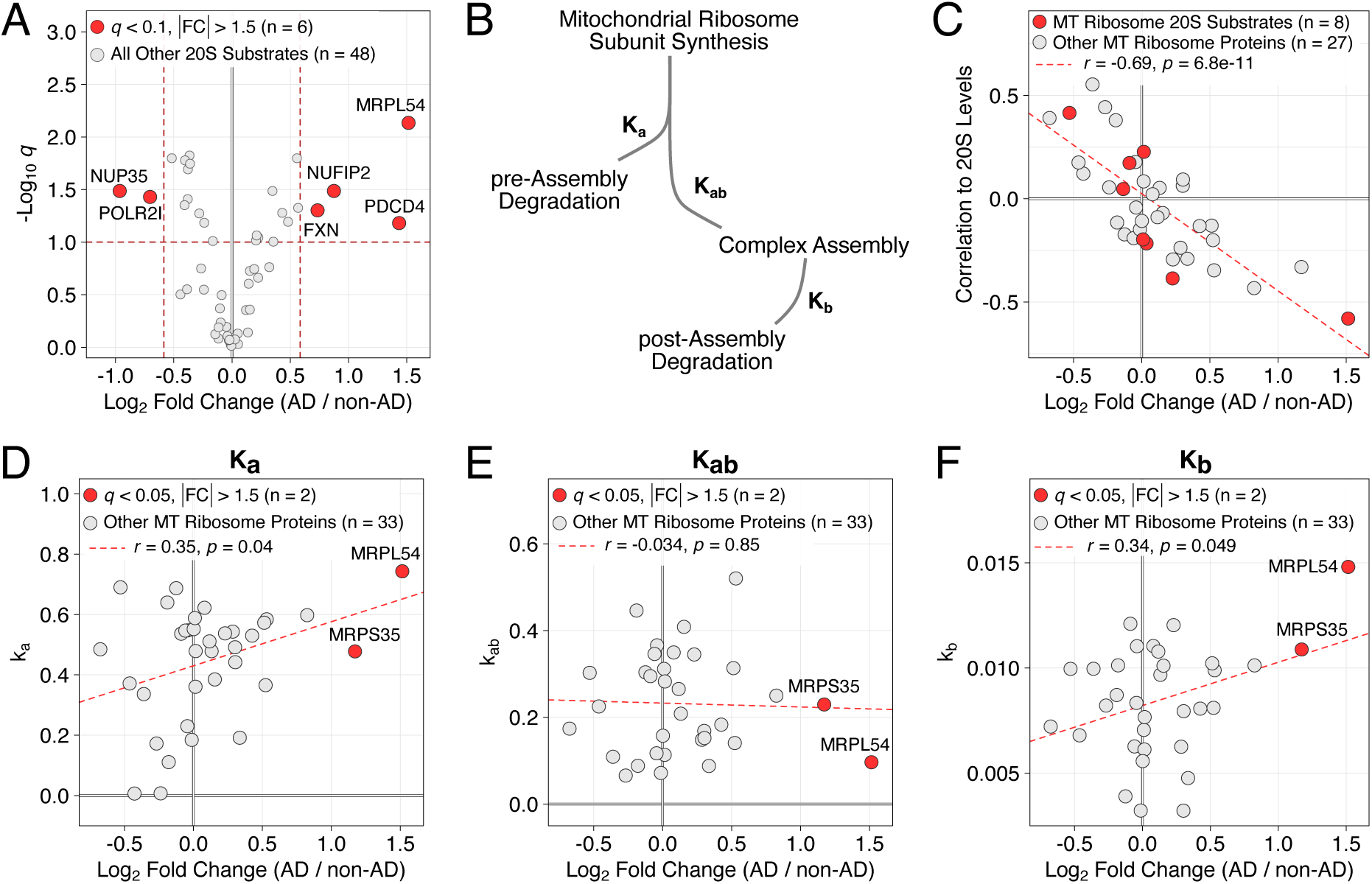
AD-associated changes in 20S substrates. **A.** Volcano plot of log_2_ fold change versus q for 20S substrates. 20S substrates are more often increased in AD. **B.** 20S correlation and log_2_ fold change are strongly negatively correlated, both among 20S substrates and all detected mitochondrial ribosome proteins. **C.** Two-step pathway of mitochondrial ribosome assembly. Based on previously-published results^109^. **D.-F.** Correlation of rate constants measuring pre-assembly degradation rate (k_a_, D.), complex assembly (k_ab_, E.), and post-assembly degradation (k_b_, F.). Mitochondrial ribosomal protein abundance is significantly correlated with both degradation rates, but not complex assembly. Note that the pre-assembly degradation rates are much higher than post-assembly (compare y-axis in D. / E.)

Based on the large and significant increase in MRPL54, we further examined the set of mitochondrial ribosome proteins in our data. Doing so revealed a highly significant relationship between the correlation of mitochondrial ribosome subunits to 20S levels and their fold change in AD (Figure 4B). Mitochondrial ribosomes are assembled through a multi-step process that requires translocating subunits encoded by the nuclear genome into mitochondria and assembling them into functional ribosomes through a series of assembly intermediates^109–111^. Select individual subunits are produced in stoichiometric excess^109,110^ and degraded with kinetics similar to those observed for other protein complexes (Figure 4C)^112^. Specifically, free subunits are rapidly degraded unless they are incorporated into functional mitochondrial ribosomes. The turnover of these subunits is thus reflected by three parameters, *k_a_* - the turnover of free subunits, *k_ab_*- the rate of complex formation, and *k_b_* - the turnover of subunits within mitochondrial ribosomes. In general, proteins within complexes are more stable than free subunits^112^ and, consequently, *k_a_* values are generally much higher than *k_b_*.

Analyzing the relationship between subunit turnover and abundance in AD revealed a previously-unappreciated relationship. Specifically, the log_2_ fold change (AD / non-AD) of individual mitochondrial ribosome subunits was significantly and positively correlated with the degradation rate constants *k_a_*and *k_b_*, but not the rate of complex formation, *k_ab_*(Figure 4D-F). That is, proteins that are rapidly degraded tended to be elevated in AD, consistent with a model in which defects in protein quality control pathways contribute to the accumulation of mis-localized, dissociated subunits of protein complexes. Notably, MRPL54, the subunit with the largest increase in AD (and among the largest increases across all proteins), has the most rapid degradation rates of both the free subunit and in complex (Figure 4D, F). Prior proteomic profiling studies have identified increases in individual mitochondrial ribosome subunits^18,19,21^ but neither they nor our study have observed systematic shifts in mitochondrial ribosomes. Our results suggest that this is, in part, because subunits that would normally rapidly be turned over preferentially accumulate, while those with slower turnover are less affected. More generally, they are consistent with a model in which proteins that are normally rapidly turned over when they mis-localize or fail to assemble into a protein complex rise to high abundance in AD.

### UPS Substrates are Increased in AD

Proteasome abundance is regulated according to the proteostatic needs of the cell^113,114^ and imbalanced proteasome subunit stoichiometry may further exacerbate AD-linked proteostasis defects^115^. We observed that 20S core particle abundance is decreased and both 19S and 20S subunits show reduced stoichiometry in AD. These observations raise the possibility that some proteins that would otherwise be targeted and degraded by the ubiquitin-proteasome system (UPS) are not efficiently removed from cells. To systematically investigate this possibility, we examined the properties of the set of differentially abundant proteins between AD and non-AD samples. We first examined the degradation rates of differentially abundant proteins. We reasoned that proteins exhibiting increased abundance owing to aberrantly diminished UPS activity would show large AD / non-AD fold changes and rapid degradation rates. Using a recently published dataset that measured degradation rates in mouse brain tissue^116^, we plotted the degradation rate and fold change, separating proteins based on fold change direction (Figure 5A). Among proteins increased in AD, we observed a significant positive correlation (*r* = 0.24, *p* = 0.024) (Figure 5A). This trend was more evident when the data were binned by log_2_ fold change, with a clear increase in degradation rate observed for proteins with a fold change greater than 1.5 (Figure 5B). In contrast, there was no relationship between degradation rate and log_2_ fold change for proteins that were decreased in AD (Figure 5A / B). These differences led us to explore relationships among differentially abundant proteins and properties of UPS substrates.

**Figure 5:**
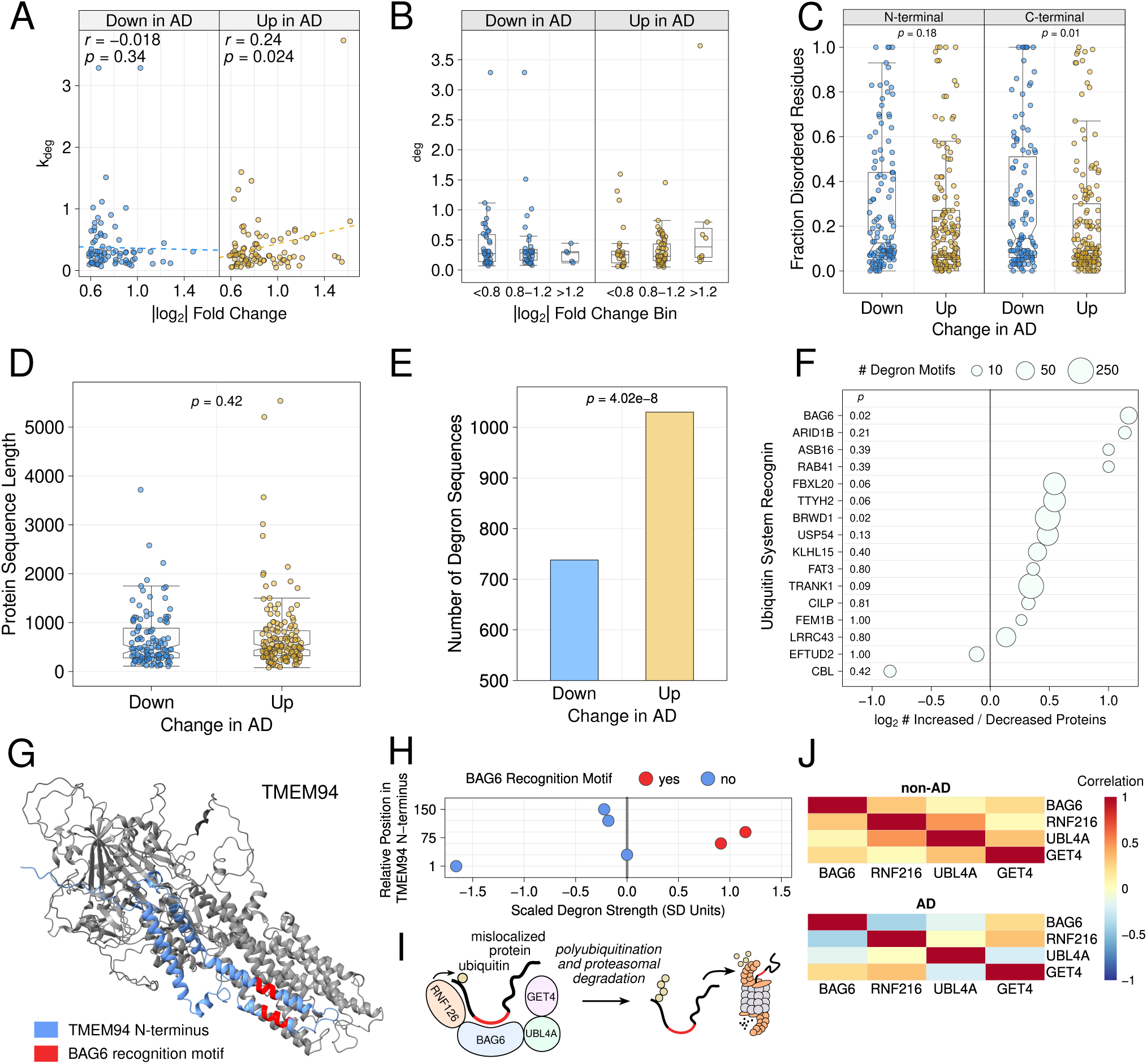
Properties of differentially abundant proteins between AD and non-AD samples. **A**. The log_2_ fold change versus degradation rate was visualized for all differentially abundant proteins stratified by the fold change direction. **B**. Binning proteins by fold change revealed a subset of proteins that had large increases in AD and high degradation rates (rightmost box). **C**. Proteins of increased versus decreased abundance do not differ in the fraction of N-terminal disordered residues. Proteins decreased in AD have a significantly higher fraction of disordered C-terminal residues. Termini were defined as the first (N-terminus) or last (C-terminus) 100 amino acids of a protein. **D**. Proteins of increased versus decreased abundance in AD have similar lengths. **E**. Proteins of increased abundance have significantly more degron-containing peptide sequences than those decreased in AD. **F**. The log_2_ ratio of the number of degron motifs in proteins of increased vs. decreased in abundance was stratified by E3 ligase. The largest and most significant difference was the enrichment of Bag6 degrons in proteins increased in AD (top row). **G**. The structure of TMEM94, a protein significantly increased in AD samples, is shown with its cytosol-facing N-terminus and Bag6 degron recognition motifs highlighted. **H**. The degron strength of peptides from the TMEM94 N-terminus were Z-transformed such that more potent degrons have higher values. Data are from a prior study^125^. **I**. Model depicting substrate recognition and processing the components of the Bag6 ubiquitin ligase complex. **J**. Correlations among the relative abundance of Bag6 complex subunits were plotted, revealing a significant decrease in subunit stoichiometry in AD samples.

Intrinsically disordered regions can facilitate the degradation of cellular proteins, potentially by acting as unstructured initiation regions for degradation^66,117,118^ or encoding ubiquitin system or proteasome recognition motifs^119,120^. We therefore explored whether proteins increased or decreased in AD differed significantly in the fraction of disordered residues in their N- and C-termini. We defined the Nand C-termini of proteins as the first or last 100 amino acids, respectively and used DISOPRED for disorder predictions^121^. At the N-terminus, differentially abundant proteins between AD and non-AD had similar fractions of disordered residues (Wilcoxon *p* = 0.18; Figure 5C). At the C-terminus, however, proteins decreased in AD had a significantly greater fraction of disordered residues, contrary to our hypothesis (Wilcoxon *p* = 0.01; Figure 5C).

We next assessed whether differentially abundant proteins differed in the number of signal sequences that allow them to be targeted for UPS degradation. The canonical pathway for UPS protein degradation involves E3 ubiquitin ligases binding degrons, then ubiquitinating substrate proteins^90^. Poly-ubiquitinated proteins are then bound and degraded by the proteasome. We searched for degrons among our differentially abundant proteins using a curated database containing thousands of human protein degron sequences^119,122–127^. We note that this database does not include modification-dependent degrons, such as the recognition of phosphorylated tau by CHIP-HSC70^128^. Despite similar overall lengths (Wilcoxon *p* = 0.42), proteins increased in AD harbored significantly more UPS degrons (Wilcoxon *p* = 4.2e-08; Figure 5D / E). These results provide further evidence that proteins that would normally be targeted for degradation by the UPS accumulate in AD.

Specific E3 ligases often target distinct classes of proteins. We next assessed whether degrons specific to individual E3 ligases were enriched in the set of differentially abundant proteins we identified. To do so, we computed the log_2_ ratio of the number of proteins of increased versus decreased abundance for degrons bound by individual E3 ligases. The results showed a clear and significant enrichment of BAG6 motifs among proteins of increased abundance. BAG6 is a chaperone protein that functions in complex with the E3 ligase RNF126, GET4, and UBL4A^129–131^. The BAG6 complex specifically identifies mis-localized proteins within the cytoplasm and participates in endoplasmic reticulum-associated degradation^129,131,132^. Increased abundance of proteins with BAG6 motifs in AD may therefore result from inadequate protein quality control-based targeting and degradation, similar to our results for mitochondrial ribosome proteins.

We examined the set of proteins of increased abundance that also contained BAG6 motifs to identify those that may accumulate due to impaired quality control mechanisms. Across all proteins identified in our study, one of the largest fold-change increases was seen for TMEM94, an ER-resident transport protein. TMEM94 contains two BAG6 motifs in its cytoplasmic-facing N-terminus (Figure 5G). To determine whether these motifs could influence TMEM94 degradation, we examined a recent proteome-wide degron scanning dataset^125^. As compared to other peptides in the TMEM94 N-terminus, the BAG6 motif peptides were strongly destabilizing, suggesting they are authentic BAG6 degrons (Figure 5H).

The BAG6 complex functions through the association of BAG6, RNF126, GET4, and UBL4^131–133^ (Figure 5I). To understand whether imbalances in the subunits of the complex might also contribute to increases TMEM94 levels in AD, we measured the correlation of individual subunits of the BAG6 complex. In non-AD samples, we observed significant positive correlations among individual BAG6 complex members. In contrast, in AD, the correlation among BAG6 subunits was reduced and, for some members, negative. Together, our results reveal a large increase in TMEM94 in AD that may result from the combination of proteasome dysregulation, as well as reduced stoichiometry of the BAG6 complex that likely targets mislocalized TMEM94 for proteasomal degradation through TMEM94’s N-terminal degrons.

### Low Correspondence of Proteomic and Transcriptomic Changes in AD

Our data suggest that impaired protein clearance, a post-translational alteration, may drive proteomic changes in AD. We hypothesized such a scenario would result in low concordance between protein versus mRNA fold changes for the same gene products. To test this hypothesis, we performed bulk RNA-seq from Brodmann area 9 of the dorsolateral prefrontal cortex for all cohort samples. We quantified the abundance of 12,890 transcripts. Of those, 488 met our significance criteria of a *q* value ≤ 0.05 and an absolute fold change difference ≥ 1.5 (Figure 6A). Multiple transcripts with well-defined roles in AD were among those increased in AD, including the neuroinflammation-related transforming growth factor-beta 1 (TGFB1), NFKB2, and nuclear factor of activated T-cells (NFATC4), as well as the parathyroid hormone receptor 1, previously linked to the physiological response to AD pathology^134^ (Figure 6A). Transcripts decreased in AD commonly reflected neuronal loss, including decreases in the cytoskeletal component TUBB2A, the calcium ion channel TRPC1, and HERC1, an E3 ubiquitin ligase that regulates synaptic membrane dynamics^135^ (Figure 6A).

**Figure 6:**
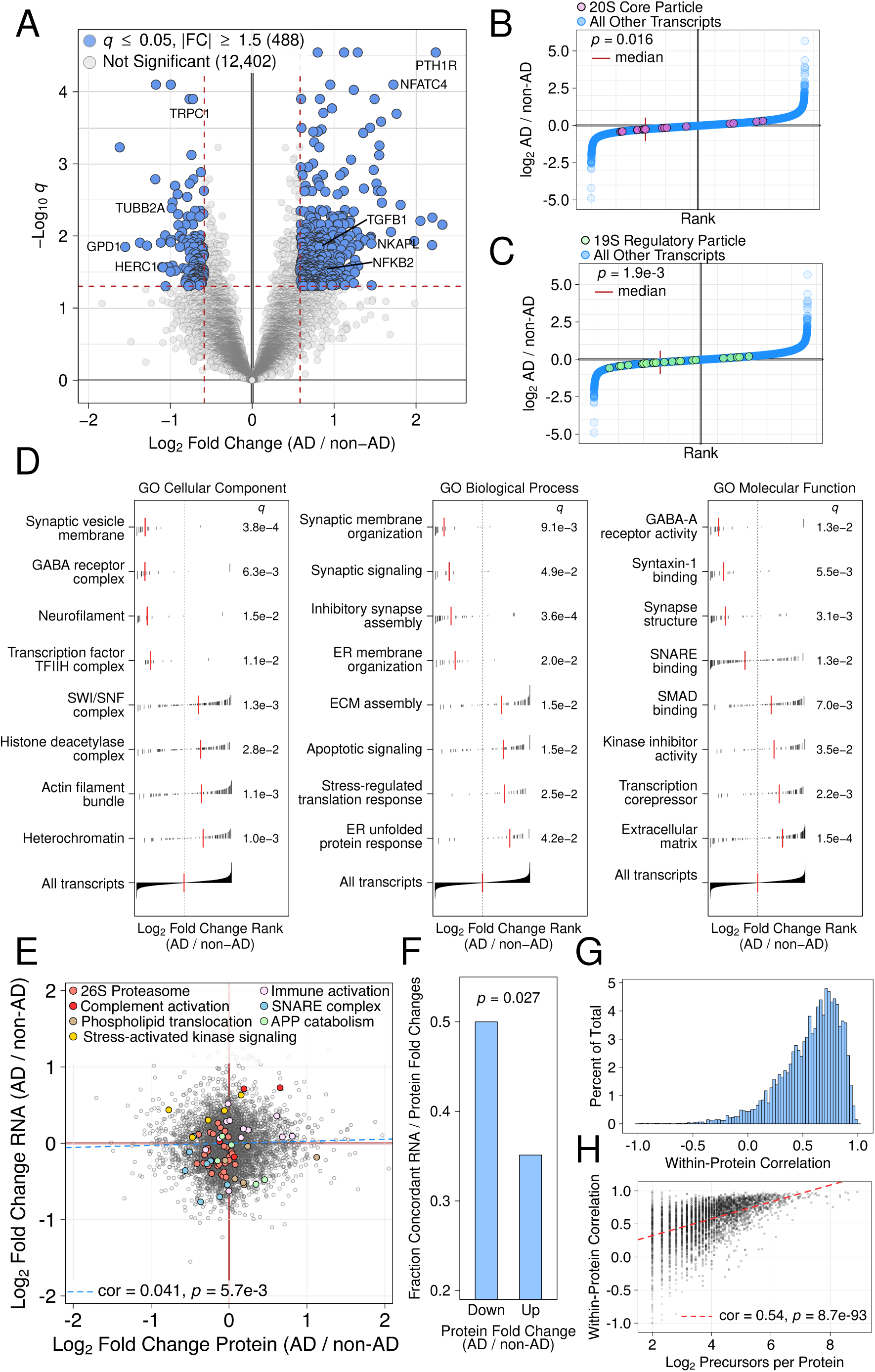
Transcriptomic changes in AD and comparison to proteomic changes. **A**. Volcano plot displaying log_2_ fold change (AD / non-AD) for all detected transcripts (12,890). Selected AD-linked transcripts are highlighted. **B**. Rank order plot displaying the fold changes of 20S subunits relative to other transcripts. The red vertical line displays the median log_2_ fold change shift for all detected 20S subunits. The p value was computed using the Wilcoxon rank sum test. **C**. As in B., but for 19S regulatory particle subunits. **D**. Gene set enrichment analysis results for all detected transcripts. Each panel shows four down-regulated and four up-regulated Gene Ontology terms from the indicated ontology. Black vertical lines display the log_2_ fold change rank among all detected transcripts while the red vertical line displays the median for all transcripts mapping to the specified term. **E**. Comparison of log_2_ fold changes for gene products detected at the protein and RNA level (6,247). Gene ontology biological process terms were used to identify sets of gene products with concordant fold changes at the RNA and protein levels, which are displayed with large, colored points. **F**. Barplot displaying the fraction of gene products with discordant fold change directions at the protein versus RNA levels. The displayed p value was computed using the chi-square test. **G**. Within-study reliability of proteomics data was estimated using split-peptide correlation analysis. The set of peptides mapping to each protein was randomly split into two non-overlapping sets and their relative abundance values were correlated. **H**. Scatterplot displaying the relationship between the number of peptides and correlation of relative among randomly selected, non-overlapping sets of peptides.

To better understand the mechanisms that might cause the decreased proteasome levels we observe in AD, we compared the levels of proteasome subunit transcripts between AD and non-AD samples. Similar to our proteomic data for the 20S proteasome, we identified a significant decrease in 20S subunit levels (*p* = 0.016, Figure 6B). In contrast to our proteomics result, levels of 19S subunits were also significantly decreased (*p* = 1.9e-3, Figure 6C). Multiple explanations may account for this discrepancy, including our ability to detect more 19S subunits at the transcript versus protein level. Post-transcriptional mechanisms may also provide a relatively stable pool of 19S subunit proteins despite decreased transcript abundance^136,137^. Taken together, the decreased levels of proteasome subunit transcripts provides a biological mechanism for the decreased proteasome levels and subunit stoichiometries we observe in AD.

To identify pathway- and complex-level alterations at the transcript level, we performed Gene Set Enrichment Analysis on the set of GO Cellular Compartment, Biological Process, and Molecular Function terms. At the Cellular Compartment level, the largest and most significant changes were decreases in neuron-specific terms, including “Synaptic vesicle membrane”, “GABA receptor complex”, and “Neurofilament” (Figure 6D). Terms increased in AD had less consistent changes and included genes related to the cytoskeleton (“Actin filament bundle”) and the regulation of gene expression (“Heterochromatin”, “Histone deacetylase complex”, and “SWI / SNF Complex”; Figure 6D). Similar patterns among terms Biological Process terms decreased in AD, which included “Synaptic membrane organization”, “Synaptic signaling”, and “Inhibitory synapse assembly”. Upregulated biological process terms reflected multiple known AD pathological processes, including cell death (“Apoptotic signaling”) and proteotoxic stress (“Stress-regulated translation response” and “ER unfolded protein response”; Figure 6D). Similar patterns were observed for Molecular Function terms, with decreased terms reflecting neuronal biology (“GABA-A receptor activity”, “Syntaxin-1 binding”, and “Synapse structure”) and increased terms reflecting disease processes (“SMAD binding”, “Transcription corepressor”; Figure 6D).

We then compared the relative fold changes for gene products quantified at both the protein and RNA levels (6,247 gene products). Overall, the correlation among protein and RNA fold changes was low, albeit positive and statistically significant (*r* = 0.041, *p* = 5.7e-3; Figure 6E), similar to prior efforts^18^. Against this backdrop, we searched for pathways whose sets of gene products exhibited consistent changes. Using Gene Ontology Biological Process terms, we identified sets of gene products with concordant and discordant changes at the mRNA versus protein levels. Among the concordant sets, 26S proteasome subunits and members of the SNARE complex were decreased at both the protein and mRNA level, while immune and neuroinflammation related gene products, including those associated with complement activation, were increased (Figure 6E). Discordant sets increased at the protein level and decreased at the RNA level included processes related to AD pathophysiology, including APP catabolism and phospholipid translocation. Specifically, the abundance of APP-related proteins, including APP itself and APOE, were increased, despite reduced levels of the corresponding transcripts, likely reflecting AD-related pathological processes. Conversely, stress-activated kinase signaling gene products were increased at the transcript level, but decreased at the protein level(Figure 6E). These results suggest that a coordinated response to ongoing pathophysiological processes occurs at the transcriptional level, but fails to propagate to the protein level, perhaps reflecting disease-associated translation defects^138^. Collectively, our results show that the overall concordance between transcriptomic and proteomic changes in AD is low, however, sets of gene products implicated in multiple disease-relevant biological processes show consistent changes at the protein versus mRNA levels.

To further evaluate our hypothesis that some proteins rise to increased abundance in AD through impaired protein clearance, we asked whether proteins of increased versus decreased abundance were more likely to exhibit consistent fold change directions at the mRNA versus protein levels. Consistent with our hypothesis, gene products increased at the protein level were significantly more likely to have a discordant fold change at the protein level (chi-square *p* = 0.027, Figure 6F). To ensure this analysis was based on reliable estimates, we restricted the set of gene products to those with at least a 0.2 absolute fold change at the protein and RNA level. However, our conclusions do not depend on this threshold, as we obtain similar results with no cutoff or an increased threshold. Our ability to accurately detect discrepancies at the mRNA versus protein levels depends on the reliability of our protein measurements. Using a previously-described approach^139^, we estimated the reliability of our proteomics data by splitting the set of peptides mapping to each protein into two randomly generated, non-overlapping sets and correlating the resulting vectors. Similar to prior results^38^, we obtain high reliability in our data (Figure 6G). As expected, the correlation between peptide sets showed a strong, positive relationship with the number of precursors mapping to each protein (Figure 6H). Thus, the low concordance among transcripts and proteins likely reflects true biological differences, rather than technical factors, such as measurement noise^139^.

### Complex Changes in Ubiquitin System Enzymes in AD

To be proteasomally degraded, many proteins must first have polyubiquitin chains (ligands for proteasome receptors) covalently attached by ubiquitin system en-zymes^63,140,141^. To understand if ubiquitin system enzymes also contribute to disease proteopathic burden alongside decreased proteasome levels in AD, we investigated how ubiquitin system components vary in AD. The ubiquitin system comprises E1 ubiquitin activating enzymes, E2 ubiquitin conjugating enzymes, and E3 ubiquitin ligases. The human genome encodes 2 known E1s, approximately 20 E2s, and approximately 600 E3 ligases^141–143^, the latter of which target proteins with distinct degrons, structures, functions, activity states, and subcellular localizations^63,65,140,141^. Some E3 ligases primarily regulate physiological protein abundance, while others contribute to protein quality control by targeting misfolded, damaged, mislocalized proteins or unincorporated protein complex subunits^63, 65, 112, 140, 141, 143^.

As expected given their large number and functional diversity, we did not identify systematic shifts in ubiquitin system enzymes (Supplementary Figure 2A). Instead, they exhibited a continuous distribution of abundance differences between AD and non-AD samples (Supplementary Figure 2B). Of note, we did not observe altered levels of CHIP, an E3 ubiquitin ligase that targets phosphorylated tau^144^, in AD (*q* = 0.51). To identify potentially informative subsets of proteins, we correlated the levels of all ubiquitin system enzymes detected in our data. We applied hierarchical clustering to the resulting matrix of pairwise correlations, which identified 16 clusters of highly correlated ubiquitin system enzymes (Figure 7A). To evaluate the statistical significance of the identified clusters, we generated an empirical null distribution by randomly sampling ubiquitin system enzymes and computing the intra-cluster correlation. We defined statistical significance as clusters that exceeded the 99% quantile of our empirical null distribution. Using this approach, we identified 12 significant clusters (Figure 7B).

**Figure 7:**
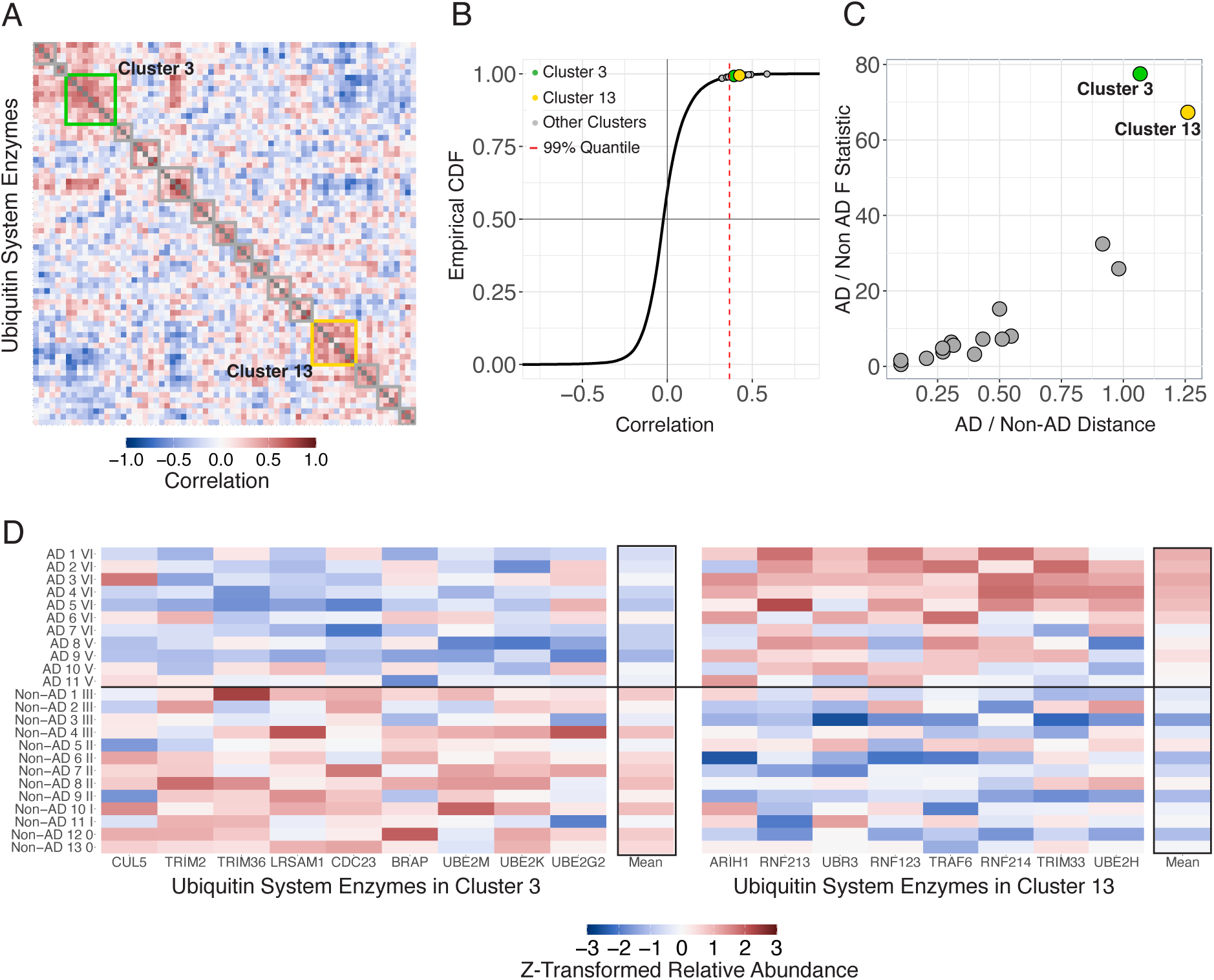
Ubiquitin system enzyme alterations in AD. **A.** Hierarchical clustering was used to identify highly correlated subsets of ubiquitin system enzymes. A total of 16 clusters, numbered from top left to bottom right were identified. **B.** To determine the statistical significance of individual clusters, we used bootstrap-based resampling to generate 1,000 random clusters. Clusters exceeding the 99th percentile of the resulting empirical null distribution were considered significant. **C.** The discriminative ability of each cluster was evaluated using two complementary methods, the scaled Euclidean distance between AD and non-AD samples (x axis) and the F statistic from an ANOVA of AD versus non-AD samples. By both metrics, clusters 3 and 13 were clear outliers. **D.** Normalized levels of all cluster 3 and 13 ubiquitin system enzymes are shown, with each cluster’s mean at right.

Our approach for defining clusters is based on protein covariation across all samples and therefore does not consider abundance differences between AD and non-AD. To determine whether individual clusters could discriminate AD from non-AD samples, we used two complementary statistics. We evaluated the scaled Euclidean distance and F statistic of an ANOVA comparing cluster levels between AD and non-AD samples. By both approaches, the third and thirteenth clusters, which contained 9 and 8 ubiquitin system enzymes, respectively, robustly separated AD and non-AD samples (Figure 7C). Strikingly, the two clusters exhibited divergent patterns, such that proteins in cluster 3 tended to show decreased abundance in AD, while those in cluster 13 showed increased abundance (Figure 7D).

Inspection of individual cluster 3 and 13 revealed multiple proteins with known or putative roles in AD. CUL5, a Cullin-RING E3 ligase, was decreased in AD (Figure 7D). The protein was recently discovered to bind and ubiquitinate tau and genetic perturbations that decrease CUL5 levels increase tau oligomer accumulation in AD models^145^. TRIM2 is a RING finger E3 ligase that binds and regulates the levels of neurofilament proteins^146^. TRIM2 is highly expressed in the central nervous system (CNS) and ablating TRIM2 expression leads to neurodegeneration in mice^146^. BRAP (BRCA1-associated protein) is also highly expressed in the CNS and reducing its levels results in aberrant histone ubiquitination patterns, as well as neurodegeneration^147^ in mice. We identified multiple E2 ubiquitin conjugating enzymes in cluster 3, including UBE2M, UBE2K, and UBE2G2 (Figure 7D). Decreased levels of selected E2s have previously been observed in AD^148,149^. Notably, both UBE2M and UBE2K are implicated in protein quality control via ERAD^150,151^, suggesting that the AD-associated decreases we observe in these proteins may further contribute to loss of protein homeostasis in AD.

Proteins in cluster 13 showed increased abundance in AD samples. TRAF6, an E3 ligase that regulates the levels of signaling proteins, has previously been shown to increase in AD^148^. Cluster 13 contained multiple RING finger E3 ligases, including RNF123, RNF213, and RNF214. Several other RING finger E3 ligases are elevated in human AD^152–154^ and our results identify additional examples of disease-associated increases in this class of E3s. ARIH1, an E3 ligase with roles in neuronal development, ubiquitinates multiple proteins with functional relevance to AD. In particular, ARIH1 modulates both microtubule stability and neurotransmitter release via its substrate targets^155,156^, suggesting that altered levels of the protein in AD may exacerbate cytoskeletal and synaptic abnormalities.

## Discussion

The full extent of protein dysfunctions in AD and the mechanisms that give rise to them have not been fully characterized. We used a recently-developed approach that provides deep proteomic coverage, highly accurate quantitation, and sample multiplexing capability^38,39^ to measure the levels of approximately 6,400 proteins in a set of 24 AD and non-AD brain tissue samples. We identified hundreds of differentially abundant proteins, including many with no previously-described role in AD, that show large increases in AD samples. We also highlight multiple mechanisms that may directly or indirectly contribute to impaired protein quality control in AD. Our analysis of protein quality control pathways and their substrates in AD reveals general principles by which proteins may aberrantly accumulate in AD-afflicted cells and exacerbate disease-linked proteotoxic stress^71,157–160^.

The largest systematic change we observed in our data was a decrease in levels of 20S proteasome subunits. This result mirrors recently described-decreases in proteasomal capacity in AD^59–61,161^. The largest decrease among individual subunits was PSMB8, a subunit of the immunoproteasome recently implicated in neurodegeneration^162^. The mechanisms that cause decreased proteasome abundance and activity in AD are not known, though multiple factors, including the accumulation of insoluble tau^163^, sequestration of the Nrf2 transcription factor that normally activates proteasome genes^59^, post-translational modifications of individual proteasome subunits^164^, and mitochondrial defects^96,145,160^ may all contribute. Our analysis of transcriptomic changes in AD suggests that the decrease in proteasome levels and activity is driven by aberrant gene expression regulation, since we observe significant and consistent downregulation of proteasome gene expression.

Unlike 20S core particle subunits, 19S regulatory particle subunits were not systematically decreased in AD at the protein level. Such a scenario could result from a decreased ratio of 20S to 26S proteasomes or imbalances in the production and assembly of 26S proteasomes. 20S proteasomes exhibit distinct substrate preferences^67,69,103,104^, in particular, targeting intrinsically disordered proteins. Multiple intrinsically disordered proteins form pathological protein aggregates in AD^70,158,165,166^, which are typically extensively ubiquitinated. Thus, a reduction in 20S proteasomes may impair the normal physiological clearance of these molecules, which may then be sequestered in aggregates as a form of molecular triage^70,99,167^. This phenomenon may be relevant to tau pathology in AD. Tau contains multiple intrisically disordered regions^168^ and in vitro, the protein is degraded by the 20S proteasome without ubiquitination^168–170^. However, phosphorylation of tau inhibits its degradation by 20S proteasomes. Thus, in AD, tau hyperphosphorylation and reduced 20S levels may result in impaired clearance that synergistically accelerates the accumulation of insoluble, aggregated tau. Tau in AD is also extensively ubiquitinated^144,171,172^, highlighting that multiple UPS pathways may target the protein for clearance in AD.

Decreased 20S subunits and unchanged 19S levels could also reflect aberrant subunit synthesis and impaired assembly, leading to the accumulation of 19S subunits. We observed a striking reduction in subunit stoichiometries of 20S, but especially 19S subunits, consistent with this notion. Prior proteomic profiling studies have described changes in proteasome subunits in AD^18,19,22,173^. Such studies typically find, similar to our results for 19S, increased levels of some subunits and decreased levels of others. Thus, aberrant proteasome subunit stoichiometry appears to be a consistent feature of AD. Proteasome assembly is a multi-step, highly regulated process that proceeds through multiple assembly intermediates with the aid of molecular chaperones^83,85,174^. Some proteasome subunits are produced in excess and unincorporated subunits are degraded through dedicated quality control pathways^83,85,112,174^. Aging, a key AD risk factor^32^, is associated with both decreased proteasome activity and stoichiometry among proteasome complex subunits, both of which likely contribute to the results observed here^80^. Our understanding of proteasome assembly’s role in aging and disease is less well-established. Although age-related defects in proteasome assembly have been described in multiple model systems^175,176^, the topic remains relatively unexplored in the context of human aging and AD.

Decreased 20S levels in AD led us to explore whether proteasome substrates accumulate in disease. We started by correlating individual proteins to 20S levels. Protein set enrichment analysis revealed multiple biological processes and cellular compartments among the set of proteins negatively correlated to 20S levels. A key theme among these terms was protein localization to a specific subcellular compartment or protein complex. This led us to explore 20S substrates exhibiting increased abundance in AD. Among these was MRPL54 which, as a component of the mitochondrial ribosome, is both specifically localized and a protein complex subunit. MRPL54 exhibited one of the largest fold increases in AD of all proteins we profiled, as well as a fast degradation rate^109^. Analysis of proteins from the mitochondrial ribosome complex revealed significant associations between subunit degradation rates and fold changes in AD, such that subunits that are normally rapidly turned over accumulate in AD. Many mitochondrial proteins are encoded in the nuclear genome^86,109,110^. They must therefore be imported into mitochondria. When the synthesis of mitochondrial proteins exceeds the translocation capacity of mitochondria, the UPS targets and degrades cytosolic mitochondrial proteins^177^. The set of 20S substrates includes mitochondrial ribosome proteins^69,104^, suggesting that decreased proteasome levels could lead to the accumulation of unincorporated subunits. More generally, our results suggest that the combination of turnover rate and subcellular localization may have utility for predicting aberrant protein accumulation in AD. In this regard, precise, proteome-wide measurements of protein turnover rates^109,116^ in AD model systems, such as induced pluripotent stem cells, would be a valuable resource.

To extend these results, we examined the properties of differentially abundant proteins in AD. Our results revealed key features of proteins that increase in abundance in AD. First, we observed a significant positive association between a protein’s turnover rate and its abundance in AD for proteins of increased, but not decreased abundance. This correlation was modest (*r* = 0.24), an expected result given that proteins may rise to increased abundance in AD through multiple mechanisms. For example, we identified increased levels of complement C4A in AD, consistent with prior studies^18,21,46^. This increase is likely driven by inflammation in AD^178^, rather than failure to properly degrade the protein. Nevertheless, we identified a significant enrichment of UPS degrons within proteins increased in AD. The most significant enrichment was a motif commonly found in mislocalized proteins that is bound by the BAG6 complex^131^. BAG6 targets mislocalized proteins, as well as aggregation prone proteins and protein fragments^129–132^. Among BAG6 degron-containing proteins, the largest increase was for TMEM94, an ER resident protein. Using data from a prior large-scale screening effort^125^, we determined that the cytosolic N-terminus of TMEM94 contains two likely authentic BAG6 degrons. TMEM94 may thus rise to increased abundance in AD both as a result of mislocalization and decreased levels and stoichiometries of the BAG6 complex, a phenomenon we also observed in AD.

Our results, therefore, are consistent with a model in which proteins rise to increased abundance in AD through the failure of normal protein clearance mechanisms. Consistent with this model, we observed low correspondence between transcriptomic and proteomic changes in disease, similar to prior results^18^. In particular, proteins of increased abundance were significantly more likely than those of decreased abundance to have discordant protein versus mRNA fold changes. Multiple lines of evidence suggest this low concordance reflects biological, rather than technical, factors. In particular, we detect many expected transcriptomic changes, such as decreases in synaptic and neuronal transcripts as well as increases in inflammation-related transcripts in AD. Likewise, analysis of our proteomic data established the reliability of our measurements, similar to those previously obtained using the same methodology^38,139^. AD is increasingly recognized as a multi-proteinopathy, with individual patients frequently exhibiting multiple co-pathologies^179^. The set of proteins known to misfold and aggregate in AD includes proteins with diverse sequences, functions and subcellular localizations, including alpha-synuclein^180^, TDP-43^8^, RBM45^181,182^, TMEM106B fragments^183^, U1-70K, and U1A^9,10^. Although misfolded proteins may be produced by diverse mechanisms in AD, their impaired clearance likely reflects more general mechanisms, such as the reduced proteosomal capacity we demonstrate here.

We also examined how levels of ubiquitin system enzymes change in AD. The ubiquitin system comprises hundreds of enzymes and we used hierarchical clustering to identify informative subsets. The levels of proteins in two clusters robustly discriminated AD from non-AD samples. Cluster 3, which contained proteins that were decreased in AD, contained Cul5. Cul5 was recently shown to ubiquitinate tau^145^ and the decrease in Cul5 we observe may further exacerbate tau dysfunction in AD. More generally, proteasome activation has long been considered a promising therapeutic target for AD^184–186^. However, our results make clear that therapeutic approaches targeting proteostasis should also consider alterations in ubiquitin system targeting of substrates.

Using plexDIA^38,39^, we quantified approximately 6,400 proteins across our 24 samples. The proteomic depth of our dataset was limited by the relatively slow scanning speed of the MS instrument used and can increase significantly by using faster instruments, such as timsTOF Ultra and Orbitrap Astral. The throughput was only 3-fold higher compared to label-free approaches since we used mTRAQ tags, which enables the simultaneous multiplexing of up to 3 samples. In principle, however, the plexDIA framework can accommodate higher plexes. Mass tags, such as the recently-developed PSMtag that can support a 9-plex, can substantially expand throughput when combined with appropriate software^187,188^. Multiplicative gains can be achieved by combining tags with recently-developed orthogonal multiplexing approaches^189^. This will support scaling the analysis to larger cohorts. Furthermore, single-cell proteomic technologies are poised to significantly increase the resolution and power of the analyses performed here^190^. Given AD’s considerable pathological heterogeneity and complex genetic and environmental risk factors^15,30–32^, the ability to profile single cells and larger cohorts would also be of great value.

Our results reveal previously-unappreciated aspects of UPS dysfunction in AD. We identify decreased levels of 20S proteasome subunits as the largest and most consistent proteomic change among cellular compartments in our data. Using protein correlations to 20S levels and publicly-available datasets, we reveal key principles of UPS substrates that increase in AD. Namely, they are rapidly turned over, they have compartment-specific subcellular localizations, they include protein complex subunits, and they are normally cleared by quality control pathways when mislocalized. Our results thus provide new insights into protein dysfunctions in AD and the mechanisms that give rise to them.

## Materials and Methods

### Cohort Selection and Tissue Samples

All postmortem frozen brain tissue samples were obtained from the Massachusetts Alzheimer’s Disease Research Center (ADRC) brain bank. A cohort of 24 patients was selected based on primary and secondary diagnoses, as well as clinical and demographic characteristics. Post-mortem neuropathological evaluations were used to classify cases on the basis of AD neuropathologic changes. Neurofibrillary tangle pathology was scored according to the Braak staging system^5^. Amyloid beta deposition was scored using the Thal staging system^191^. To increase statistical power to detect disease-associated proteomic changes, we classified Braak V or VI cases as “AD”, while all other subjects were classified as “non-AD”. Subjects were chosen so that the cohort contained similar numbers of AD and non-AD cases, as well as males and females. Table 1 provides detailed demographic information for each subject. Approximately 2 g of prefrontal prefrontal cortex tissue (Brodmann area 9) was dissected from each case and immediately stored at −80 ^◦^C until processed.

**Table 1:** Subject demographics. “ADRC” corresponds to the patient number, “PMI” - post-mortem interval (hours) between death and tissue collection, “Primary / Secondary Dx” - primary or secondary diagnosis, respectively, “CVD” - cardiovascular disease, “CAA” - cerebral amyloid angiopathy, “ARS” -atherosclerosis. Ages greater than 90 are listed as “90+” per IRB rules regarding patient privacy.

| ADRC | Age | Sex | PMI | Braak Stage | Thal Stage | Primary Dx | Secondary Dx |
| --- | --- | --- | --- | --- | --- | --- | --- |
| 1628 | 60 | F | unknown | VI | 4 | AD | CVD |
| 1636 | 86 | M | 20 | VI | 3 | AD |  |
| 1669 | 86 | M | 10 | I | 0 | control |  |
| 1703 | 73 | F | 20 | 0 | 0 | control |  |
| 1821 | 92 | M | unknown | II | 0 | control |  |
| 1837 | 68 | M | 27 | I | 0 | control |  |
| 1845 | 90+ | F | 19 | VI | 5 | AD | CVD, CA |
| 1854 | 85 | F | 6 | III | 2 | AD | CVD, CA |
| 1886 | 58 | F | 18 | 0 | 0 | control |  |
| 1906 | 71 | M | 12 | III | 4 | AD | CVD, CA |
| 1907 | 90+ | F | 14 | III | 2 | AD |  |
| 1926 | 82 | F | 6 | V | 4 | AD | CVD, CA |
| 2015 | 90+ | M | 39 | III | 3 | control | AD |
| 2018 | 90+ | F | 24 | II | 1 | control |  |
| 2068 | 79 | F | 9 | II | 0 | control | ARS |
| 2112 | 78 | M | 5 | VI | 4 | AD | CV |
| 2132 | 90+ | F | 30 | V | 4 | AD | CV |
| 2191 | 87 | M | 21 | II | 3 | control | CVD, AD |
| 2203 | 75 | F | 10 | VI | 5 | AD | CVD |
| 2223 | 90+ | M | 21 | VI | 4 | AD | CV |
| 2225 | 90+ | F | 14 | V | 4 | AD | CAA |
| 2232 | 65 | F | 11 | V | 5 | AD | CVD |
| 2233 | 72 | F | 10 | VI | 5 | AD | CVD, ARS |
| 2259 | 90+ | F | 30 | II | 3 | AD | CVD |

### Bulk Brain Tissue Processing and Protein / RNA Extraction

Bulk tissue samples were processed to generate cell lysates using bead-based tissue disruption, as previously described^192^. Approximately 50 mg of tissue was transferred to a microcentrifuge tube containing zirconium oxide beads (Next Advance 430917) and 600 *µ*l lysis buffer on ice. The lysis buffer contained 75 M NaCl, 50 mM EPPS (pH 8.5), 10 mM sodium pyrophosphate, 10 mM sodium orthovanadate, 3% SDS, 10 mM PMSF, and one EDTA-free protease inhibitor tablet (Roche 11873580001). After adding the tissue, the tubes were transferred to a Mini-Beadbeater 16 (Biospec). Samples were processed for 30 seconds then placed on ice for 2 minutes. This process was then repeated three times to ensure complete tissue lysis.

We used the single-pot, solid phase-enhanced sample preparation (SP3) method^193^ to extract and purify peptides from brain tissue lysates. The SP3 workflow uses paramagnetic beads that bind proteins via hydrophilic interactions to separate proteins from complex mixtures^193^. Equal amounts of Sera-Mag E3 and E7 beads (5 mg; Cytiva 65152105050250 [E3] and 45152105050250 [E7]) were added to a microcentrifuge tube. To condition the beads, the tube was placed in a magnetic rack, the supernatant was removed, 200 *µ*l of mass spectrometry grade water was added, and the mixture was gently mixed by pipetting after removal from the magnetic rack. This process was repeated twice. The conditioned beads were then added at a 10:1 (wt:wt) ratio to the tissue lysates and mixed by gentle pipetting. One volume of 100% ethanol was then added to the bead-lysate mixture to induce protein binding to the beads. To enhance protein binding to beads, tubes were incubated in a thermal mixer at 24 ^◦^C for 5 minutes with shaking at 1000 rpm. The tubes were then returned to the magnetic rack and the supernatant removed. The tube was then removed from the magnetic rack and washed three times with 180 *µ*l of 80% ethanol. The samples were then air dried. We carried out on-bead tryptic digestion of proteins to peptides by re-hydrating the samples in 100 mM triethylammonium bicarbonate buffer (TEAB; pH 8.5) and trypsin. Tubes were gently inverted to mix the beads and solution. Each tube was subsequently sonicated in a water bath to disrupt bead aggregates. Digests were carried out overnight by incubating samples on a thermal mixer set to 37 ^◦^C for 18 hours with shaking at 1000 rpm. Peptide abundances were quantified by absorbance at 280 nm using a NanoDrop Eight spectrophotometer (Thermo). Peptide supernatants were then transferred to new tubes and evaporated to dryness. Samples were resuspended in 100 mM TEAB.

To multiplex the analysis of brain tissue samples, peptide digests were labeled with mTRAQ using a previously-described approach^38^. We used the Δ0, Δ4, and Δ8 tags (Sciex 4440015, 4427700, and 4427698, respectively) for sample labeling. Patients were assigned to batches, where each batch contains one sample each tagged with Δ0, Δ4, and Δ8 tags. Samples were assigned to batches using block randomization so that each batch had a similar distribution of age, sex, and disease stage. Each mTRAQ tag was resuspended in isopropanol, then added a concentration of 0.1 U per 10 *µ*g of peptides per sample. The tag labeling reaction was carried out by incubating the samples at room temperature for 2 hours. Labeling reactions were quenched by adding 0.25 % hydroxylamine to the samples and incubating for 1 hour at room temperature, as previously described^38^. After quenching, samples were pooled based on the batching scheme described above.

Total RNA was extracted from all samples using the Monarch Total RNA miniprep kit (New England Biolabs T2010S) according to the manufacturer’s instructions. All samples were processed simultaneously as a single batch. Briefly, we cut and dissociated approximately 10 mg of tissue from each case and incubated the samples in the supplied proteinase K solution. We vortexed the samples, transferred them to collection tubes, then washed and eluted RNA in nuclease-free water. RNA concentration and purity were assessed using a Nanodrop 8 (Thermo). RNA samples were stored at −80 ^◦^C until being processed for RNA-seq.

### LC-MS Analysis

Sample batches were analysed by LC-MS using an Orbitrap Exploris 480 MS (Thermo) coupled to a Vanquish Neo LC system (Thermo). For each sample batch, 1 *µ*g of peptides was loaded onto an Aurora Ultimate C18 (IonOpticks AUR3-25075C18; 25 cm x 75 *µ*m) column. Samples were separated using a 135 minute gradient consisting of varying amounts of 0.1% formic acid in MS-grade water (buffer A) and 80% acetonitrile (ACN), 0.1% formic acid in MS-grade water (buffer B). The gradient started at 5% buffer B, increased to 7% buffer B within 0.5 minutes, then ramped to 32% buffer B over 120 minutes, and finally increased to 95% buffer B over the final 2 minutes. The column was washed at 95% B for 8 min, before dropping back to 5% in 0.1 min. The flow was kept constant at 200 nL / min. The total MS acquisition time per sample was 135 min and data was acquired in data-independent acquisition (DIA) mode.

To avoid contaminating the instrument with excess labeling reagent at the beginning of the gradient, the electrospray voltage was off during the first 5 minutes of each run and only set to 1900 V at minute 5. Since a droplet accumulates at the end of the emitter tip, an in-house developed assembly was used to blow it off at the sweep gas outlet on the source and used for a time-dependent flow of 5 arbitrary units of sweep gas between minutes 4.5 and 5 of the method duration, as previously described^194^. The temperature of the ion transfer tube was 275 ^◦^C. One duty cycle consisted of 2x (1 MS1 scan and 30 MS2 scans). The MS1 scans were conducted in profile mode at a resolution of 120K with a scan range from 378 - 1372 m / z with RF lens level of 50% and a normalized AGC target of 300%. The first round of MS2 scans spanned a mass range of 380 - 620.5 m/z. The DIA windows were 8.5 Th wide with 0.5 Th overlaps. The normalized collision energy was set to 30, the orbitrap resolution was 30K, the RF lens level was set to 50%, the normalized AGC target was set to 1000%, and the maximum injection time was set to auto mode. The second round of MS2 scans was conducted at the same settings, but the mass range was 620 - 1370.5 m/z with 8.5 Th width for the first 8 DIA windows, then 17.5 Th for 9 windows, then 41.5 Th for 13 windows. The MS2 scans were acquired in centroid mode.

In a separate experiment, we used gas phase fractionation of a pooled sample of brain tissue lysate protein digest labeled with mTRAQ Δ0 to create an empirical spectral library for searching our raw MS data^44^. To do so, we first pooled equal amounts of peptides from three samples (ADRC numbers 1845, 2097, and 2225). We then labeled the pooled sample with mTRAQ Δ0 using the labeling protocol described above. We injected 1 *µ*g of labeled, pooled sample and six fractions were collected in triplicate. Library fractions were then analyzed on Orbitrap Exploris 480 MS coupled to a Vanquish Neo LC system. Library fractions were profiled using the buffers and acquisition settings described above with the following modifications: 500 ng of peptide were loaded onto the C18 column. The gradient started at 4% B, increased to 5% B within 0.5% min, then ramped to 28% B over 120 min, and finally to 95% B over 2 min. The column was washed at 95% B for 8 min, before dropping back to 4% in 0.1 min, with a constant flow rate of 200 nL/min. The MS1 scans were conducted in profile mode at a resolution of 120K with RF lens level of 50% and a normalized AGC target of 300%. The m / z ranges were: 379 - 501 m / z, 499 - 621 m / z, 619 - 741 m / z, 739 - 861 m / z, 850 - 1101 m / z, 1099 - 1341 m / z. For the MS2 scans, the normalized collision energy was set to 30, the orbitrap resolution was 60K, the RF lens level was set to 50%, the normalized AGC target was set to 1000% and the maximum injection time was set to auto mode. The MS2 scans were acquired in centroid mode. The first four fractions used 2.5 Th windows with 0.5 Th overlap, while the last two used 4.5 Th windows with 0.5 Th overlap.

### LC-MS Data Processing

#### Library Generation

Raw files from our gas phase fractionated samples were used to create an empirical library using DIA-NN (version 1.9.2)^41^. First, an in silico library was created using FASTA files of AD-relevant proteins from literature and the mTRAQ label was added as described previously^38^. The resulting predicted library was then used to search the replicate injections of each fraction to generate empirical libraries, which were then combined into one final library.

#### Protein Identification

We first searched the gas phase fractionated spectra to create an empirical spectral library using DIA-NN (version 1.9.2)^41^. The generated library was used to search all samples. We used the following DIA-NN settings to search the gas phase fractionated data: N-terminal methionine excision: enabled, peptide length: 7 - 30 amino acids, precursor m / z: 375 - 880, and charge: 1 - 5, variable modifications: methionine acetlyation and N-terminal oxidation, and fixed modifications: mTRAQ (N-terminal, K). Channel-specific normalization was used and mass accuracy was set to 10 ppm for both MS1 and MS2. The match-between-runs feature was enabled.

### RNA-seq

Total RNA samples obtained from Brodmann area 9 were shipped on dry ice to Novogene for RNA-seq analysis. Samples were subject to three quality control assays to measure sample concentration, purity, and integrity. Total RNA concentration and sample purity were assessed using the Nanodrop UV-Vis spec-trophotoemeter (Thermo). Sample integrity was assessed using the Tapestation gel electrophoresis system (Agilent). All samples were sent for sequencing after quality control. Libraries were prepared for RNA-seq and sequenced on a NovaSeq Plus (Illumina) instrument with paired-end 150 bp reads and targeting an average depth of 20 million reads per sample.

### Data Processing and Analysis

#### Raw Proteomics Data Processing and Statistical Analysis

Raw data was processed using a modified version of a previously-described approach^38^. To improve quantitative accuracy, we first corrected for potential isotopic envelope interference^38^. To do so, we computed the theoretical distribution of isotopes for each precursor and used this distribution to correct the signal from mTRAQ labels for MS1-level signal. We used the ‘diann_maxlfq’ function from the DIA-NN R package^41^ (which implements the LFQ algorithm) to quantify protein group abundances based on MS1 area. Quantified protein groups were subsequently batch corrected for mTRAQ label and run using ComBat^195^. Batch correction and all subsequent statistical analyses were carried out in the R statistical computing environment^196^.

For the statistical analysis of the batch corrected, protein-level data, we first centered the columns of the proteins (rows), patient (columns) matrix to the median protein abundance for each patient. To make the quantification relative, we then normalized each protein to the mean of the 24 samples. We tested for differential abundance between AD and non-AD using a parametric F-test^195^, followed by false discovery rate (“FDR”) correction using the Benjamini-Hochberg method. We considered proteins significantly differentially abundant when they had an FDR level *q* value ≤ 0.05 and an absolute fold change greater than 1.5 between AD and non-AD.

For the analysis of cellular compartment shifts, we used a previously-described approach for detecting systematic shifts in proteins annotated to a specific cellular compartment^56^. Briefly, we used the “MSigDB”^197^ R package, which queries the Molecular Signatures Database^198,199^ to map proteins in our dataset to GO cellular compartment terms, then tested whether the terms are significantly shifted using a Wilcoxon test. Pairwise correlation matrices for various protein complexes, including the proteasome 20S core particle and 19S regulatory particle were computed as Pearson correlations. Protein set enrichment analyses were carried out using the Wilcoxon rank sum test for the set of proteins mapping to each Gene Ontology term, similar to prior studies^139^. For all protein set enrichment analyses, we considered GO terms significant at a 0.05 FDR.

We used several publicly available tools and datasets for additional analyses. We used the “UniProt.ws” R package (DOI: 10.18129/B9.bioc.UniProt.ws) to download protein sequences and DISOPRED^121^ to predict the fraction of terminal disordered residues in each protein. For the analysis of UPS degrons in differentially abundant proteins, we used a curated database of E3 ubiquitin ligase-degron pairings and searched for matching sequences in our data. The TMEM94 structure was obtained from Alphafold^200^ and visualized using ChimeraX^201,202^. Data on protein degradation data in mouse brain samples, mitochondrial ribosome turnover and assembly, and the degron potency of N-terminal TMEM94 peptides sequences were obtained from publicly available datasets from previous works^109,116,125^.

To identify informative sets of ubiquitin system enzymes, we first correlated the relative level of each pair of E3 Ligases across all samples in our batch corrected matrix, *r_ij_* for all pairs of proteins *i* ≠ *j*. As these correlations represent a measure of similarity, we can obtain a natural dissimilarity matrix as *d_ij_* = 1 − *r_ij_*. We formed clusters using the complete linkage approach, whereby each pair of proteins is initialized into distinct, individual clusters. Then, the two clusters with minimum dissimilarity (or maximum correlation) are merged together. Next, for each iteration, the maximum dissimilarity is computed between each cluster, as the maximum distance across all pairs of members between two clusters. The clusters associated with the smallest maximum dissimilarity are merged together, and this process continues until all points have been merged into a single cluster. In order to obtain distinct clusters, we cut the hierarchical tree based on height. Branches connected above this height are considered members of the same cluster. As the tree is computed using 1− correlation as the dissimilarity, and correlation can be measured as −1 to 1, the tree has maximum height 2, and clusters are formed in this case by cutting at 1.

To provide a measure of the statistical significance of each cluster, we generated an empirical null distribution by sampling random clusters of ubiquitin system enzymes. We randomly permuted cluster labels 1000 times and at each iteration, we calculate the average pairwise correlation among members of permuted clusters. We compare the average pairwise correlation among members of our selected clusters to that of the bootstrap clusters. Note that, as performing hclust results in clustering for the entire set of pairwise protein correlations, the distribution of permuted cluster correlations reflects a local, and not global, null distribution. We considered significant clusters to be those exceeding the 99% quantile of the empirical null distribution.

To evaluate the ability of ubiquitin system enzyme clusters to distinguish AD from non-AD samples, we used two complementary metrics. We first computed the F statistic by applying an ANOVA model to ubiquitin system enzymes present in each cluster, with AD status as the only explanatory variable. We used Euclidean distance, a complementary measure of discriminative ability, to further evaluate clusters. To obtain a scaled Euclidean distance, we computed the average relative protein intensity across patients with the same disease status for each ubiquitin system enzyme in each cluster. Next, we computed the euclidean distance between averages across disease status, and divided by the number of proteins in each cluster so as to ensure comparable distances between clusters of different sizes.

#### Raw RNA-seq Data Processing and Statistical Analysis

Raw RNA-seq reads were processed as follows. We used fastQC^203^ to provide an initial overview of the sequencing quality. The resulting reports were used to assess per base sequence quality, per sequence quality scores, per base sequence content, per sequence GC content, sequence duplication levels, and adapter content. We aggregated fastQC reports into a single, interactive report using multiQC with default settings^204^. To remove adapters and low-quality bases, we used fastp^205^ with default settings. We quantified transcript abundance using Salmon^206^. To minimize spurious mapping of reads, we first constructed a decoy-aware transcript index. To do so, we downloaded human transcriptome and genome sequences from Gen-code (gencode.v38.transcripts.fa.gz and GRCh38.primary_assembly.genome.fa.gz, respectively) then built the index using the ‘salmon index’ command. We then quantified transcript abundance using this index and the ‘salmon quant’ command. To detect differentially abundant transcripts, we used salmon quant output files with DESeq2 with disease status (AD versus non-AD) as the sole contrast. As in our analysis of proteomic data, we considered differentially abundant transcripts to be those with a *q* (corrected *p*) value ≤ 0.05 and an absolute fold change ≥ 1.5. Gene set enrichment analysis was performed using the set of Gene Ontology terms similar to the proteomic data and prior studies^139^.

## Data Availability

Raw proteomic data used for all analyses described in the manuscript have been deposited to the ProteomeXchange consortium via the PRIDE partner repository with the accession number: PXD064265.

Raw and processed RNA-seq data has been deposited in the Gene Expression Omnibus and is available at the following accession number: GSE327021

The code used for data analysis and to produce figures is available at: github.com/mac230/AD_bulk_plexDIA

## Acknowledgments

PTI is a Convergent Research Focused Research Organization (FRO) and has received support from Eric and Wendy Schmidt as well as Griffin Catalyst. The Massachusetts ADRC is supported by NIH grant P30AG062421. We thank the members of PTI for constructive feedback on the project and manuscript and Andrew Leduc of Northeastern University for the mouse brain protein turnover data.

## Competing Interest

N.S. is a founding director and CEO of Parallel Squared Technology Institute, which is a nonprofit research institute. The authors declare no other competing interest.

## Author Contributions

*Experimental design*: NS, BH, MC, AP, CF

*Sample preparation and data acquisition*: PS, TCS, AM, KA

*Raising funding and supervision*: NS, AP, BH, DO, MC

*Data analysis*: ME, CF, JD, MC

*Writing and editing*: MC, JD, CF, ME, BH, NS

## Supplementary Materials

**Supplementary Figure 1:**
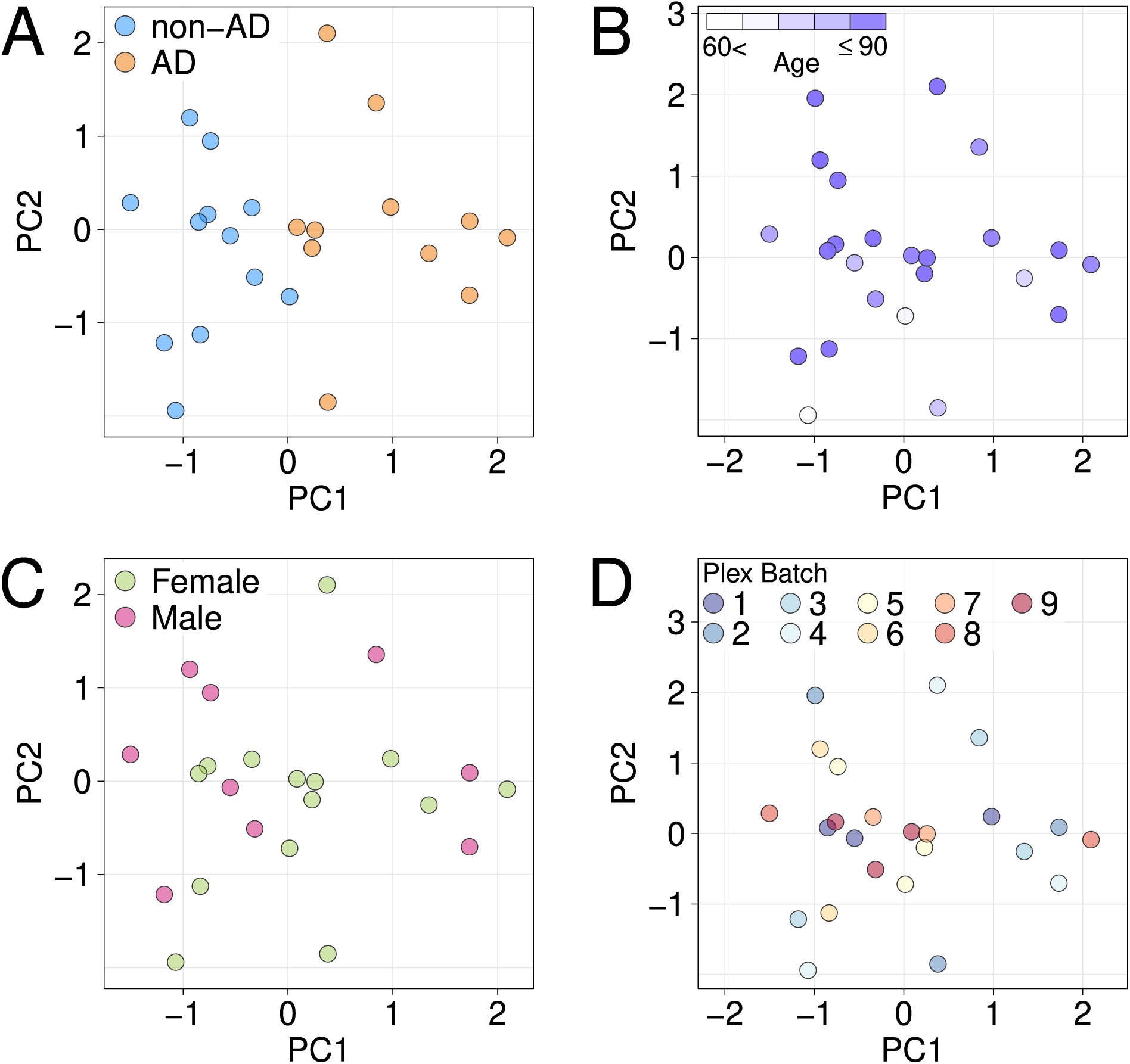
Principal components analysis (PCA). PCA was used to visualize sample similarity across relevant biological and technical variables. The plots show samples plotted along the first and second principal components and colored according to **A.** AD status, **B.** binned age, **C.** sex, and **D.** sample batch.

**Supplementary Figure 2:**
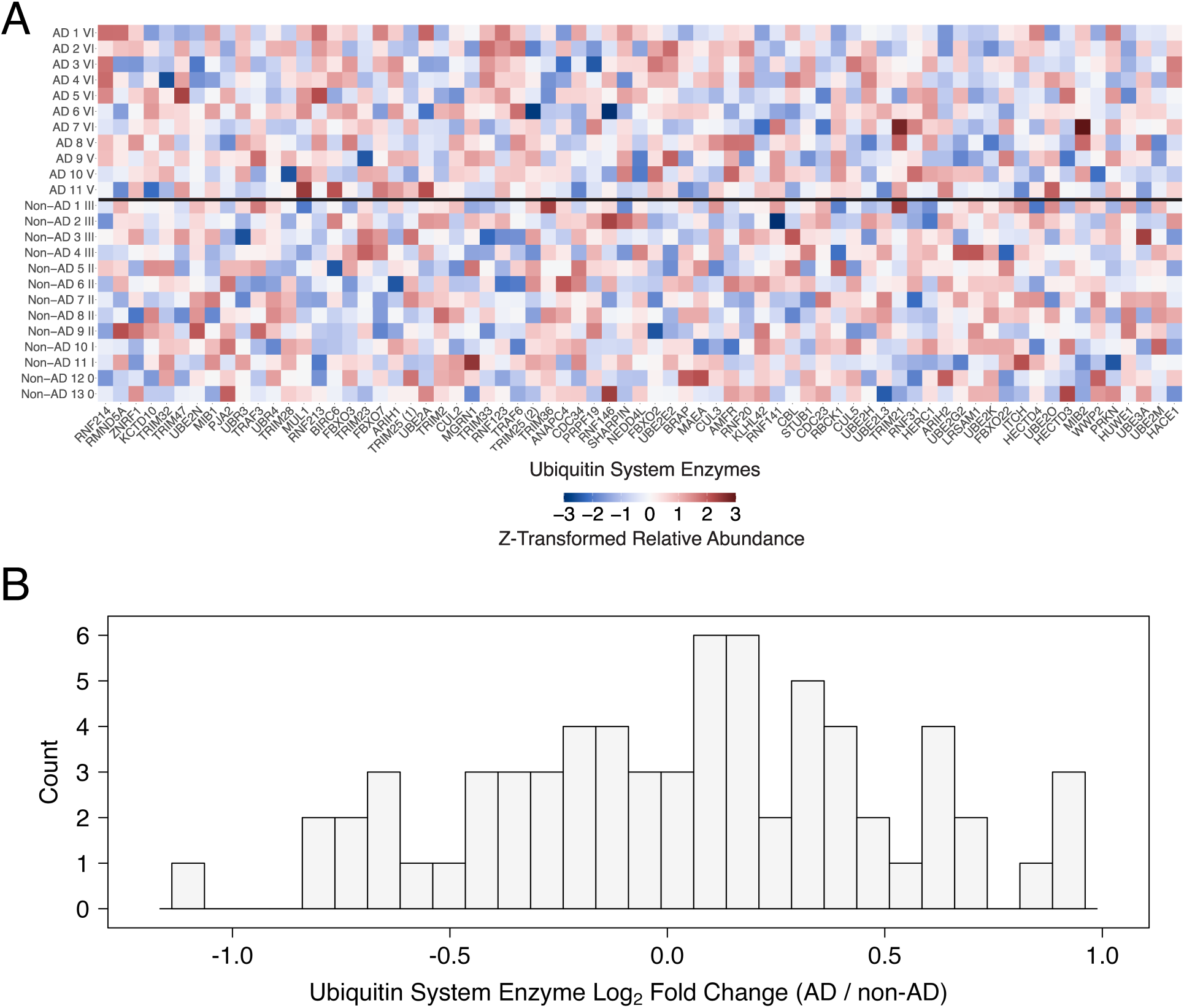
Heatmap of ubiquitin system enzymes colored by relative abundance for all 24 samples. Individual ubiquitin system enzymes are not clustered and samples are separated according to disease status (AD or non-AD), highlighting an absence of systematic shifts in the abundance of ubiquitin system enzymes.

## Notes

https://www.ebi.ac.uk/pride/archive/projects/PXD064265

https://www.ncbi.nlm.nih.gov/geo/query/acc.cgi?acc=GSE327021

### Summary of Updates

This version of the manuscript has been revised to include additional analyses that demonstrate that the decrease in proteasome levels is the largest systematic shift in protein complexes in AD detected in our data. New analyses based on RNA-seq data are also included to provide additional evidence for the notion that many proteins rise to increased abundance because of aberrant decreases in protein turnover.

